# Regulation of defective mitochondrial DNA accumulation and transmission in *C. elegans* by the programmed cell death and aging pathways

**DOI:** 10.1101/2021.10.27.466108

**Authors:** Sagen E. Flowers, Rushali Kothari, Yamila N. Torres Cleuren, Melissa R. Alcorn, Chee Kiang Ewe, Geneva Alok, Pradeep M. Joshi, Joel H. Rothman

**Affiliations:** Department of MCD Biology and Neuroscience Research Institute, University of California Santa Barbara, CA, USA; Computational Biology Unit, Institute for Informatics, University of Bergen, Norway

**Keywords:** heteroplasmy, uaDf5, purifying selection, programmed cell death, aging, insulin signaling

## Abstract

The heteroplasmic state of eukaryotic cells allows for cryptic accumulation of defective mitochondrial genomes (mtDNA). “Purifying selection” mechanisms operate to remove such dysfunctional mtDNAs. We found that activators of programmed cell death (PCD), including the CED-3 and CSP-1 caspases, the BH3-only protein CED-13, and PCD corpse engulfment factors, are required in *C. elegans* to attenuate germline abundance of a 3.1 kb mtDNA deletion mutation, *uaDf5*, which is normally stably maintained in heteroplasmy with wildtype mtDNA. In contrast, removal of CED-4/Apaf1 or a mutation in the CED-4-interacting prodomain of CED-3, do not increase accumulation of the defective mtDNA, suggesting induction of a non-canonical germline PCD mechanism or non-apoptotic action of the CED-13/caspase axis. We also found that the abundance of germline mtDNA*^uaDf5^* reproducibly increases with age of the mothers. This effect is transmitted to the offspring of mothers, with only partial intergenerational removal of the defective mtDNA. In mutants with elevated mtDNA*^uaDf5^* levels, this removal is enhanced in older mothers, suggesting an age-dependent mechanism of mtDNA quality control. Indeed, we found that both steady-state and age-dependent accumulation rates of *uaDf5* are markedly decreased in long-lived, and increased in short-lived, mutants. These findings reveal that regulators of both PCD and the aging program are required for germline mtDNA quality control and its intergenerational transmission.

## Introduction

Mitochondrial diseases are a group of conditions that affect mitochondrial functions in up to 1 in 4,300 people [1–3]. Generally, these diseases present as dysfunction in the tissues or organs with the most intensive energy demands, most commonly in muscle and the nervous system [4]. Many of these diseases are attributable to mutations in the mitochondrial DNA (mtDNA) or nuclear DNA (nDNA), and include those disorders with defects in mitochondrial function, dynamics, or quality control, or in which there is miscommunication between the mitochondria and the endoplasmic reticulum (ER) [3, 5]. The progressive advancement of the diseased state resulting from age-dependent accumulation of mutant mtDNA is a common trait among mitochondrial diseases [6–8]. While the severity of the disease varies with the nature of the mutation, the most severe phenotypes result in childhood death, as in Leigh syndrome and MELAS [5, 9]. As there are currently no pharmacological treatments for mitochondrial diseases, it is of great importance to uncover the cellular processes that underlie the regulation of mtDNA quality control.

mtDNAs show high mutation rates [10–12] and hence it is critical that cells possess mechanisms to remove detrimental mtDNA alleles, a process called purifying selection [13, 14]. Defects in this process can result in mitochondrial diseases, allowing harmful mtDNA mutations to persist through the maternal germline and subsequent generations. Processes that regulate mtDNA quality control include mitochondrial fission/fusion dynamics and mitophagy [8,15–20], and the mitochondrial unfolded protein response (UPR^MT^; [21–27]. Further, it has also been found that the insulin/IGF-1 signaling pathway (IIS; [28, 29]) ameliorates the fitness defects of mutant mtDNA.

One potential cellular process that could be used to eliminate defective mtDNAs is the culling of cells bearing mtDNA mutations by programmed cell death (PCD). The mechanisms of both developmentally controlled and genotoxicity-induced PCD have been shown to be well-conserved across metazoans [30], and much of the machinery that choreographs this process is directed by mitochondria [31–33]. Mitochondrial-dependent processes participating in PCD include permeabilization of the inner mitochondrial membrane and release of mitochondrial factors that mediate transduction of intermediary events in the cell suicide program [31–33]. Both mitochondrial function and PCD are linked to the process of organismal aging [34]. mtDNA mutations accumulate in tissues as organisms age, and it has been suggested that this accumulation is a major contributor to aging [35–37].

The nematode *C. elegans* provides an attractive model for exploring the potential role of PCD in mtDNA purifying selection. The well-described, conserved PCD regulatory pathway in *C. elegans* functions not only to eliminate 131 somatic cells during development through a rigidly stereotyped program [38], but is also activated apparently stochastically during germline development, resulting in the death of >95% of nuclei that would otherwise be destined to become oocytes in the mature hermaphrodite [30,39–41]. In addition to this “physiological” PCD, germline nuclei that have experienced genotoxic stress are eliminated through p53-dependent apoptosis, as is also the case in somatic mammalian cells [42, 43]. Thus, germline PCD allows for selective removal of nuclei with damaged genomes, thereby preventing intergenerational transmission of defective nuclear DNA. Given the prominent role played by mitochondria in the PCD process, it is conceivable that mitochondrial dysfunction could trigger PCD in the germline and, as such, might similarly provide a quality control mechanism for eliminating aberrant mtDNA, as seen for the nuclear genome.

We report here that germline mtDNA quality control in *C. elegans* is influenced by regulators of both PCD and the aging program. We find that pro-apoptotic regulators of germline PCD, notably the caspases CED-3 and CSP-1, the BH3-only domain protein CED-13, and regulators of cell corpse engulfment, reduce abundance of an mtDNA deletion and that abrogation of their functions results in elevated levels of the defective mtDNA. Notably, however, loss of the CED-3 activator CED-4/Apaf1 [44, 45] does not result in elevated levels of defective mtDNA. These findings raise the possibilities that either the caspases and the other pro-apoptotic factors function in mtDNA purifying selection by a non-canonical CED-4-independent cell death program, or that these pro-apoptotic regulators function in mtDNA purifying selection through a PCD-independent mechanism. We also report that defective mtDNA accumulates in the germline of animals with age and that although the abundance of the defective mtDNA is reduced in offspring, progeny of older mothers inherit higher levels of the mutant mtDNA than those from young mothers. Intergenerational removal of the defective mtDNA appears to be enhanced in older animals with defective mtDNA quality control. Further, we found that lifespan-extending mutations in both the IIS pathway and the non-IIS-dependent lifespan-regulator CLK-1/MCLK1 decrease accumulation of defective mtDNA, and that short-lived mutants show elevated accumulation, implicating molecular regulators of the aging process in mtDNA purifying selection. Our findings reveal that the PCD machinery and the aging program contribute to the removal of mtDNA mutations during germline development and their intergenerational transmission.

## Results

### The stably maintained mtDNA deletion mutant *uaDf5* contains multiple linked mutations resulting in aberrant proteins and shows deleterious effects on growth

To test the role of potential regulatory factors in mtDNA purifying selection, we took advantage of *uaDf5*, a 3.1 kb mtDNA deletion mutation that removes part or all of four protein-coding genes and seven tRNAs (Fig. 1A) [46]. Given the presumably deleterious nature of this defective mtDNA, it was of interest to understand how it is stably transmitted despite active purifying selection processes. While its maintenance at high levels is attributable in part to stabilization by the mitochondrial UPR [25], *uaDf5* persists, albeit at lower levels, in animals lacking this activity. One possible explanation for this phenomenon is that the mutant mtDNA might be maintained in heteroplasmy with an otherwise intact mtDNA carrying a complementing mutation. We sought to test this possibility through deep sequencing of mtDNA isolated from the *uaDf5*-bearing strain. Comprehensive sequence analysis revealed that, in addition to the large deletion, the strain indeed carries a second mtDNA mutation, *w47* (Fig. 1A, Fig. 1 – figure supplement 1A, B, C). *w47* is a single base pair insertion in the *nduo-4* gene that causes a frameshift, predicted to result in a truncated NADH dehydrogenase 4 (ND4) protein lacking 321 residues (Fig. 1 – figure supplement 1D). ND4 is an essential transmembrane subunit within complex I of the mitochondrial respiratory chain (MRC), which drives NADH-oxidation-dependent transport of protons across the inner mitochondrial membrane [47–49]. While this raised the possibility of two complementing mtDNA genomes, we found that the *w47* mutation is present at the same abundance as the *uaDf5* deletion mutation (∼75% of total mtDNA, Fig. 1 – figure supplement 1B) rather than that of the wild-type mtDNA, strongly suggesting that it resides on the same mtDNA genome. As this second mutation cannot explain stabilization of the defective mtDNA by *trans*-complementation of two deleterious mutations, other mechanisms appear to promote the stable inheritance of *uaDf5*.

**Figure 1:**
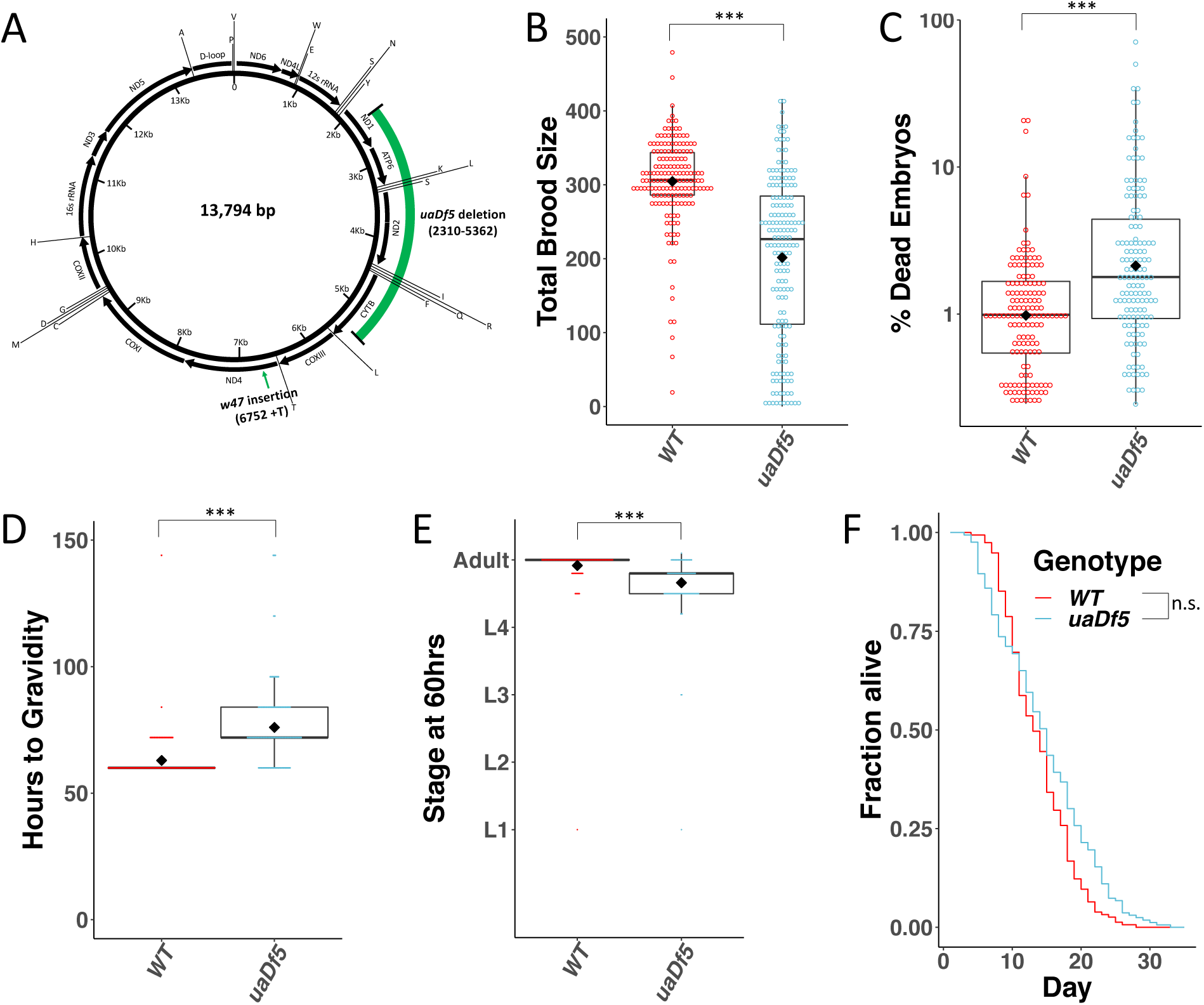
Analysis of the impact of mtDNA*^uaDf5^* on fitness parameters. **(A)** Diagram of *C. elegans* mtDNA. Black bars with arrows indicate the locations of genes and direction of transcription. Black lines with letters indicate the locations of tRNAs. Green bars show the locations of the mtDNA*^uaDf5^* deletion as well as the linked *w47* insertion that was identified via Illumina sequencing. **(B)** Brood size analysis of mtDNA*^uaDf5^* compared to laboratory wildtype N2. **(C)** Embryonic lethality analysis of *uaDf5* compared to laboratory wildtype N2. **(D)** Developmental rate analysis of mtDNA*^uaDf5^* compared to laboratory wildtype N2, counting how many hours it takes for starved L1s to reach gravidity once plated on food. **(E)** Developmental rate analysis of mtDNA*^uaDf5^* compared to laboratory wildtype N2, staging worms 60 hours after synchronized, staved L1s are plated on food. **(F)** Survival curve analysis of mtDNA*^uaDf5^* compared to laboratory wildtype N2, day 1 is defined as the day starved L1s are plated on food. Median lifespan and statistics are presented in Supplemental Figure 2. For **B-E**, box plots show median and IQR, and the diamond indicates the mean. Statistical analysis was performed using the Mann-Whitney test. (*** p < 0.001, n.s. not significant).

In addition to the aberrant protein encoded by the *w47* frameshift mutation in *nduo-4*, a second abnormal protein is encoded by the *uaDf5* genome: one end of the deletion results in a fusion protein comprised of the first 185 amino acids of NADH dehydrogenase 1 (ND1, a homolog of the core MT-ND1 transmembrane subunit of complex I of the MRC [47,49,50]), and the last 81 amino acids of mitochondrial-encoded cytochrome b (CTB-1/CYTB, a transmembrane subunit of complex III of the MRC [47,49,51]) (Fig. 1 – figure supplement 1E). It is conceivable that accumulation of these two abnormal proteins -- the truncated ND4 and the ND1-CYTB fusion protein resulting from *w47* and *uaDf5*, respectively – activate the UPR^MT^, which has been shown to result in clearance of mtDNA*^uaDf5^*, dependent on the ATFS-1 transcription factor [25, 27].

Animals harboring mtDNA*^uaDf5^* are viable and fertile, presumably because they contain intact wildtype mtDNA [46]. However, we found that *uaDf5-*bearing animals displayed a significant reduction in brood size (WT 304 ± 4.8; *uaDf5* 201 ± 8.6 embryos laid, p < 0.001) (Fig. 1B) and a significant increase in embryonic lethality (WT 1.4 ± 0.2%; *uaDf5* 4.2 ± 0.7%, p < 0.001) (Fig. 1C). Additionally, *uaDf5* animals are slow-growing, evident in both the number of hours to reach gravidity (WT 63 ± 0.8; *uaDf5* 76 ± 1.4 hours at 20°C, p < 0.001) (Fig. 1D) and the stage of development reached after 60 hours of feeding (WT: adult; *uaDf5:* mid-L4) (Fig. 1E). In contrast, however, we were surprised to find that lifespan was not substantially affected (WT 14 ± 0.4; *uaDf5* 15 ± 0.5 days) (Fig. 1F, Fig. 1 – figure supplement 2). Given the significant decline in the majority of fitness parameters tested, we conclude that *uaDf5* is a useful tool for studying mitochondrial disease and mechanisms underlying mtDNA quality control.

### PCD regulators promote removal of mtDNA*^uaDf5^*

During germline development in *C. elegans*, as many as 95% of nuclei destined to become potential oocytes are eliminated by PCD [39–41,52]. While this process has been proposed to be stochastically determined [39, 52], it has also been suggested that it may function to selectively remove all but the most “fit” germline cells. As such, PCD could perform a role in purifying selection in the germline, wherein potential oocytes that undergo PCD are associated with higher levels of defective mtDNA. To test this hypothesis, we introduced *uaDf5* into various PCD mutants and quantified abundance of the defective mtDNA by digital-droplet PCR (ddPCR; see Supplementary Table 1 for list of mutants tested). In a wildtype genetic background, we found that the steady-state fractional abundance of *uaDf5* in populations of 200 day 1 adults (first day of adulthood) is highly reproducible across four separate trials, demonstrating the reliability and robustness of the assay. Our analyses confirmed that mtDNA*^uaDf5^* constitutes the major molar fraction of mtDNA in the *uaDf5*-bearing strain by a nearly 3:1 ratio (Fig. 2 – figure supplement 1).

CED-3 in *C. elegans* is the major executioner caspase in the canonical PCD pathway [38,52,53] and is required for virtually all PCD both in the germline and the soma (Fig. 2 – figure supplement 2). We found that two *ced-3* mutations that strongly block PCD [54] showed a significant increase in the ratio of defective to normal mtDNA from a molar ratio of 2.9:1 for *ced- 3(+)* to 3.5:1 for *ced-3(n717)* and 4.6:1 for *ced-3(n1286)* (Fig. 2A). This effect appears to be exclusively attributable to an increase in mtDNA*^uaDf5^* in the PCD-deficient strains, as the abundance of mtDNA^WT^ is not significantly different between the strains. These *ced-3(-)* mutations both localize to the p15 domain of the protease portion of CED-3 (Fig. 2 – figure supplement 3), consistent with abolition of caspase activity. These findings implicate the CED-3 caspase and its p15 domain in mtDNA quality control. We found that one other mutation located in the p15 domain showed only a very slight increase that was not statistically significant (*n2454:* 2.9:1, Fig. 2B). While it is unclear why this allele showed a weaker effect it is noteworthy that, unlike the other two mutations, which result in a dramatic alteration of the protein, this mutation is predicted to result in a relatively modest (ala → thr) single amino acid substitution.

**Figure 2:**
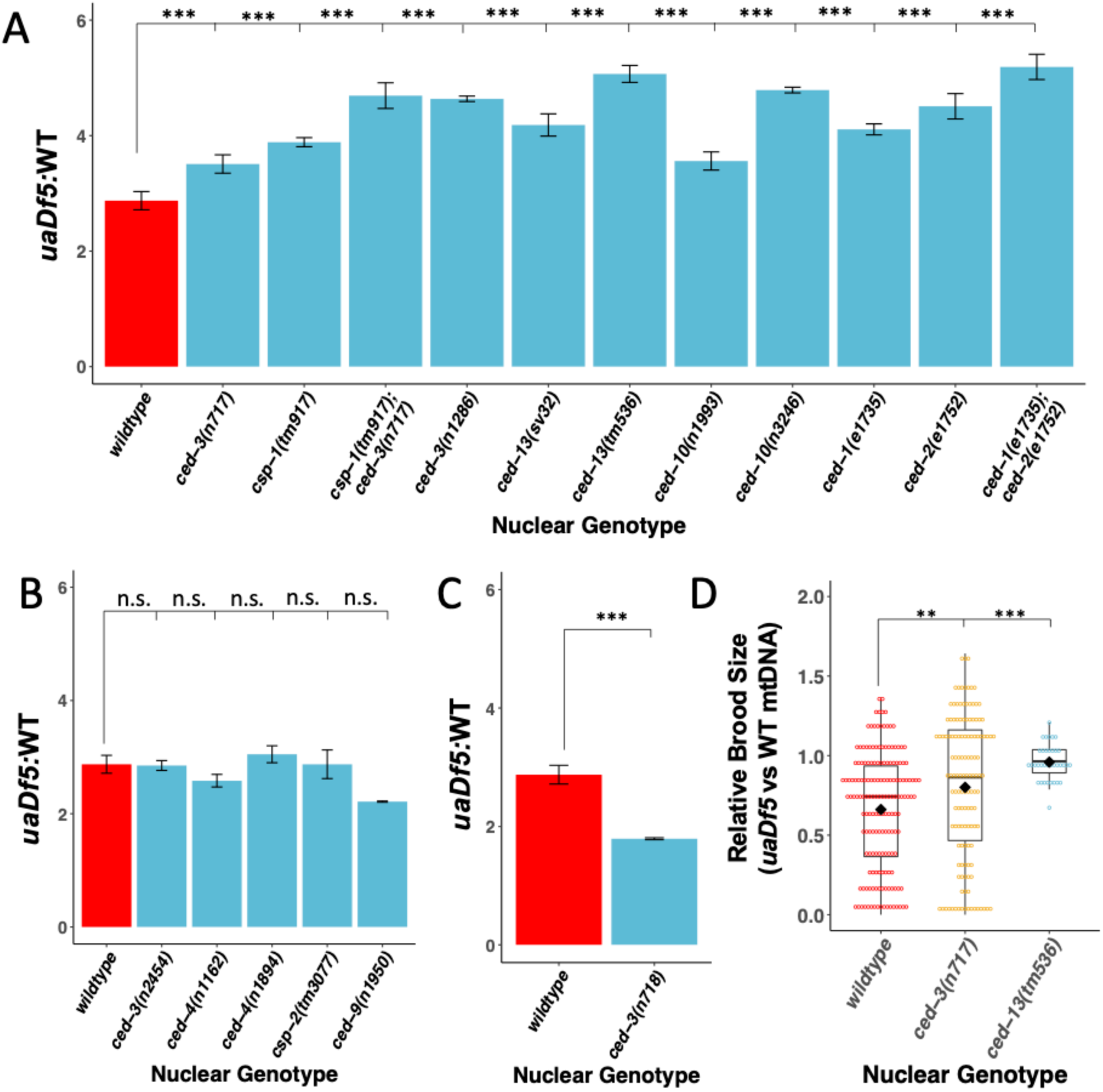
Regulators of PCD act on mutant mtDNA. **(A-C)** ddPCR analysis of the steady-state molar ratio of mtDNA*^uaDf5^* in 200 worm populations of day 1 adults of various PCD mutant backgrounds. **(A)** PCD mutants that result in a significant increase in the molar ratio of mtDNA*^uaDf5^*. **(B)** PCD mutants that result in no statistical change in the molar ratio of mtDNA*^uaDf5^*. **(C)** PCD mutant that results in a significant decrease in the molar ratio of mtDNA*^uaDf5^*. **(D)** The relative brood size of the animals with and without mtDNA*^uaDf5^* in the indicated mutant backgrounds. For each nuclear genotype shown, the brood size of *uaDf5*-containing worms was normalized by dividing by the average brood size of worms containing only *WT-mtDNA*. Box plots show the median and IQR, the diamond indicates the mean. For **A-C**, n=3 or more biological replicates of 200 worm populations were performed for each genotype. Statistical analysis was performed using one-way ANOVA with Dunnett’s correction for multiple comparisons. For **D**, statistical analysis was performed using the Mann-Whitney test. Error bars represent SEM. (*** p < 0.001, ** p < 0.01, * p < 0.05, . p < 0.1, n.s. not significant).

A second caspase in *C. elegans*, CSP-1, also functions, albeit less prominently, in PCD. While loss of CSP-1 alone does not result in a strong reduction in PCD, it synergizes with loss of CED-3 both in PCD and in other caspase-dependent processes (Fig. 2 – figure supplement 2) [55–57]. We found that removing CSP-1 in the *csp-1(tm917)* knockout mutant results in a significant increase in mtDNA*^uaDf5^* abundance to a molar ratio of 3.9:1 (Fig. 2A). Further, we found that this mutation enhances the effect of the *ced-3(n717)* mutation, increasing the mtDNA*^uaDf5^*:mtDNA^WT^ molar ratio from 3.5:1 to 4.7:1 (Fig. 2A). Together these findings demonstrate that caspase activity, and possible PCD-mediated clearance, are crucial for mtDNA quality control and function in purifying selection of defective mtDNA.

We sought to further investigate a potential role for PCD in mtDNA purifying selection by evaluating the requirement for the pro-apoptotic factor CED-13, a BH3-only domain protein that acts specifically in the germline to activate PCD [38,58,59]. Consistent with a requirement for PCD in purifying selection, we found that two *ced-13* alleles result in a very substantial increase in the mtDNA*^uaDf5^*:mtDNA^WT^ molar ratio (*sv32*: 4.2:1 and *tm536*: 5.1:1) (Fig. 2A), supporting the notion that CED-13 promotes removal of defective mtDNA in the germline. CED-13 functions in PCD by antagonizing the function of mitochondrially localized CED-9/Bcl-2 [58, 59], which normally sequesters the apoptosome factor CED-4/APAF1 at mitochondria, thereby preventing it from triggering autocatalytic conversion of the executioner caspase zymogen proCED-3 to its pro-apoptotic protease structure (Fig. 2 – figure supplement 2) [60]. An equivalent action is carried out in the soma by the BH3-only protein EGL-1 [60]. *n1950*, a gain-of-function allele of *ced-9* that blocks the interaction of EGL-1 with CED-9 at the mitochondria, results in elimination of PCD in the soma but not the germline [39,52,60,61]. Consistent with the lack of effect of *ced-9(n1950gf)* on germline PCD, we found that the mtDNA*^uaDf5^*:mtDNA^WT^ molar ratio was not increased in *ced-9(n1950gf)* mutants (2.2:1) (Fig. 2B). Thus, CED-3, CSP-1, and CED-13 are required both for germline PCD and for removal of mtDNA*^uaDf5^*.

Cells that undergo PCD are cleared by the surrounding cells in the process of engulfment and degradation, which is implemented through a set of redundant pathways that converge on the CED-10 GTPase [62–65] (Fig. 2 – figure supplement 2). Although this engulfment process is necessary primarily for removal of the resultant corpses, it also appears to play an active role in cell killing: inhibition of the engulfment pathway diminishes occurrence of PCD, likely through a complex feedback mechanism [66, 67]. Further supporting a role for PCD in purifying selection, we found that single or double mutations of several genes that promote engulfment of cell corpses result in elevated mtDNA*^uaDf5^*:mtDNA^WT^ molar ratios, ranging from 3.5:1 to 5.2:1 (4.1:1 for *ced-1(e1735)*, 4.5:1 for *ced-2(e1752)*, 5.2:1 for the *ced-1(e1735); ced-2(e1752)* double mutant, 3.6:1 for *ced-10(n1993)*, and 4.8:1 for *ced-10(n3246)*; Fig. 2A).

We tested whether *generally* increased germline PCD alters mtDNA*^uaDf5^* abundance by examining the effect of removing the caspase-related factor, CSP-2, which has been shown to play an anti-apoptotic role through inhibition of CED-3 autoactivation in the germline (Fig. 2 – figure supplement 2) [68]. We found that a loss-of-function mutation in *csp-2*, which elevates germline PCD, did not alter the mtDNA*^uaDf5^*:mtDNA^WT^ molar ratio (2.9:1 for the *csp-2(tm3077)* knockout mutation) (Fig. 2B). This observation does not conflict with a requirement for PCD in purifying selection: a general increase in PCD in the germline of *csp-2(-)* animals would not be expected *per se* to alter the mechanism that *discriminates* defective from normal mtDNAs and therefore the relative abundance of the two forms. Rather, our findings suggest that the mechanisms that recognizes and disposes of the defective mtDNA specifically requires PCD components acting in a selective, rather than general process (i.e., in those cells with the highest burden of the defective mtDNA).

### Evidence for non-canonical action of PCD regulators in mtDNA purifying selection

The foregoing results implicate a role for the pro-apoptotic CED-3 and CSP-1 caspases, CED-13, and the CED-1, 2, and -10 cell corpse engulfment factors in mtDNA purifying selection. However, several observations suggest that elimination of normal, physiological germline PCD *per se* is insufficient to increase accumulation of defective mtDNA, or that these factors promote purifying selection through a PCD-independent pathway. Specifically, we observed that the occurrence of PCD does not perfectly correlate with their effects on accumulation of mtDNA*^uaDf5^* in particular mutants.

First, we found that the *ced-3*(*n718)* allele lowers, rather than elevates, the abundance of mtDNA*^uaDf5^* (mtDNA*^uaDf5^*:mtDNA^WT^ molar ratio of 1.8:1, Fig. 2C). This effect is likely to be attributable to the nature of the *n718* mutation. Those *ced-3* mutations that result in increased mtDNA*^uaDf5^* levels (Fig. 2A) both alter the p15 domain, which is essential to the active caspase function. In contrast, the *n718* mutation changes a residue in the caspase activation and recruitment domain (CARD), located within the prodomain of the CED-3 zymogen, which is removed upon caspase activation and affects its activation by CED-4 (Fig. 2 – figure supplement 3) [44, 54]. While *ced-3*(*n718)* strongly compromises PCD, this mutation might not alter CED-3 caspase function in a way that interferes with its role in mtDNA purifying selection.

Our surprising finding that while CED-3 activity is required for mtDNA purifying selection, a CED-3 mutation that compromises its activation by CED-4 did not elevate mtDNA*^uaDf5^* levels prompted us to investigate the requirement of CED-4 in mitochondrial purifying selection. Consistent with the effect of the *ced-3*(*n718)* mutation, we found that eliminating the function of the pro-apoptotic regulator CED-4, the *C. elegans* orthologue of mammalian Apaf1 and the upstream activator of CED-3 in the canonical PCD pathway [60, 69], did not result in a marked increase in the relative abundance of mtDNA*^uaDf5^*. That is, while the mtDNA*^uaDf5^*:mtDNA^WT^ molar ratio increased to 3.1:1 in the *ced-4(n1894)* mutant, the effect was not statistically significant. Moreover, the canonical allele *ced-4(n1162)* allele similarly showed no elevation in mtDNA*^uaDf5^* (molar ratio = 2.6:1; Fig. 2B). These results suggest that CED-3 caspase functions in mitochondrial purifying selection independently of the caspase-activating factor CED-4.

### Evidence that decreased fitness, but not lifespan, is attributable to mtDNA*^uaDf5^*-induced PCD

Taken together, our results implicate many PCD regulatory factors, and potentially PCD, in the selective clearance of defective germline mtDNAs. Our additional observations suggest that defective mtDNAs may, in fact, *trigger* elevated germline PCD, resulting in the production of fewer mature gametes and progeny. Specifically, we found that the significant decrease in brood size that we observed in *uaDf5-*bearing animals with a wildtype nuclear background is partially suppressed by both *ced-3(-)* and *ced-13(-)* mutations, which prevent PCD (Fig. 2D, Fig. 2 – figure supplement 4A), suggesting that elimination of PCD might rescue cells that would otherwise be provoked to die as a result of accumulation of defective mtDNA. Our findings further underscore the observation that accumulation of defective mtDNA in those animals that do survive does not affect longevity, as we found that lifespan is unaltered in these PCD mutants even when the levels of mtDNA*^uaDf5^* are nearly doubled (Fig. 2 – figure supplement 4B, C).

### Age-dependent accumulation of mtDNA*^uaDf5^* in the germline

Our findings that PCD regulators are required to reduce mtDNA*^uaDf5^* abundance, the central role that mitochondria play in PCD [31–33,70], the observed decline of mitochondrial health during the aging process [7,35–37,71–75], and the relationship between excessive PCD and the aging phenotype [34] led us to examine the dynamics of mtDNA*^uaDf5^* accumulation as worms age. We measured the fractional abundance of *uaDf5* in adults at progressively increased ages spanning day 1, defined as the first day of egg-laying, through day 10. Day 1 through day 4 of adulthood encompasses the time during which nearly all self-progeny are produced. After day 4, hermaphrodite sperm become depleted and the animals transition into a post-gravid, progressively aging state [76–78]. By day 10, animals exhibit indications of advanced age. Analysis of the abundance of mtDNA*^uaDf5^* revealed a progressive increase throughout gravidity and post-reproductive aging, with the mtDNA*^uaDf5^*:mtDNA^WT^ molar ratio increasing from 2.9:1 to 5.5:1 (Fig. 3A, Fig. 3 – figure supplement 1A). This age-related accumulation of *uaDf5* in adult worms is reminiscent of the accumulation of mtDNA mutations seen in aging mammals [35,73,79] and suggests that *uaDf5* in *C. elegans* may be a useful tool for studying the role that mtDNA mutations play in aging.

**Figure 3:**
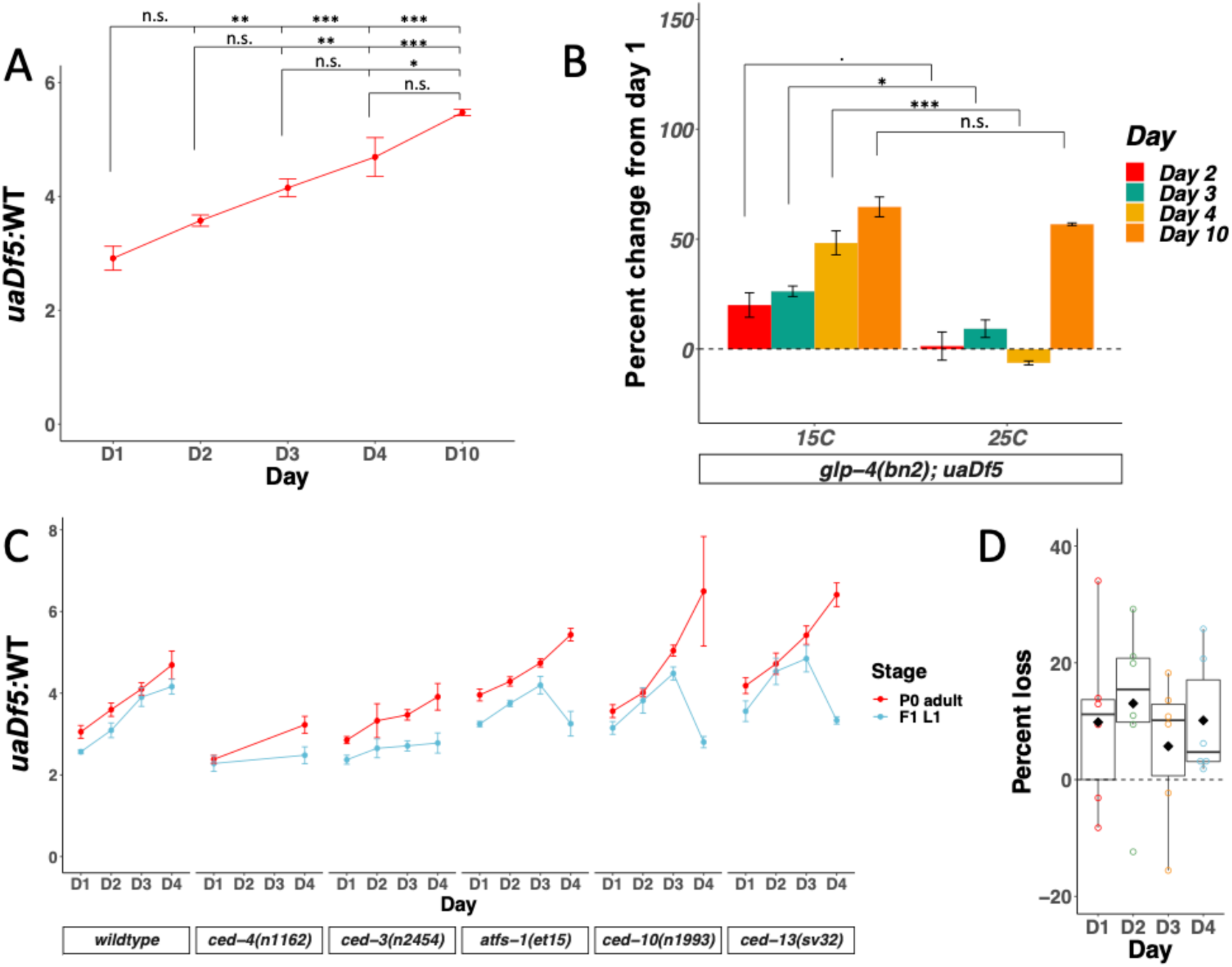
mtDNA*^uaDf5^* accumulates in the germline of aging adults, and evidence of purifying selection between mother and offspring. **(A)** Analysis of the molar ratio of mtDNA*^uaDf5^* in aging adults in a wildtype nuclear background. **(B)** Analysis of the percent change of *uaDf5:*WT from day 1 (Y axis = (*uaDf5:*WT day x -*uaDf5:*WT day 1)/(*uaDf5:*WT day 1)). For *glp-1(q231ts), fem-3(q20ts),* and *glp-4(bn2ts),* 15°C is the permissive temperature (germline development occurs) and 25°C is the restrictive temperature (female germline development is inhibited). Statistical analysis was performed using one-way ANOVA with Tukey correction for multiple comparisons. (*** p < 0.001, ** p < 0.01, * p < 0.05, . p < 0.1). Error bars represent SEM. **(C)** Analysis of the molar ratio of mtDNA*^uaDf5^* in aging adults (P0 adult) and their L1 progeny (F1 L1) in various nuclear backgrounds shows that all strains decrease the *uaDf5* load during transmission from mother to offspring, and that strains with significantly higher mtDNA*^uaDf5^* levels (*atfs-1(et15), ced-10(n1993)*, and *ced-13(sv32)*) have a more significant removal mechanism at day 4 of adulthood. n=3 or more replicates of 200 worm populations were performed for each timepoint. Error bars represent SEM. Grey dashed line indicates a hypothetical threshold at which high mtDNA*^uaDf5^* burden activates enhanced intergenerational purifying selection in older mothers. **(D)** Analysis of the measured loss of mtDNA*^uaDf5^* between mother and offspring at each day of adulthood shows that mtDNA*^uaDf5^* removal occurs. n=6 replicates of 200 worm populations for each condition.

Given that gametes are depleted with age, it is conceivable that the age-dependent increase in mtDNA*^uaDf5^* is attributable to accumulation in somatic mitochondria. To assess whether the observed age-related accumulation of mtDNA*^uaDf5^* occurs predominantly in the maternal germline or in somatic cells, we analyzed animals defective in germline development by taking advantage of the *glp-4(bn2)* mutant, which produces only a small number (∼12) of germline cells compared to that in wildtype animals (∼1,500), with no known effect on somatic gonad development [80]. In contrast to the increased mtDNA*^uaDf5^* abundance with age seen at permissive temperature (mtDNA*^uaDf5^*:mtDNA^WT^ of 1.8:1 at day 1, rising to 2.6:1 at day 4, for an overall increase by 48%), we found that *glp-4(bn2)* animals at the non-permissive temperature showed a slight decrease in the defective mtDNA from day 1 to day 4 of adulthood (2.2:1 at day 1, dropping to 2:1 at day 4, for an overall decrease of 6.3%) (Fig. 3B). These results strongly suggest that the observed age-dependent increase in mtDNA*^uaDf5^* abundance occurs exclusively in the germline. We found that mtDNA*^uaDf5^* does eventually appear to accumulate in somatic cells with age, as day 10 adults grown at the restrictive temperature showed a marked increase in the mtDNA*^uaDf5^*:mtDNA^WT^ molar ratio compared to day 1 adults even in the absence of a germline (day 10 *glp-4(bn2)* molar ratio of 3.4:1, a 57% increase from day 1 levels) (Fig. 3B). We conclude that the marked increase in mtDNA*^uaDf5^* with age during the period of fecundity occurs primarily in the germline and that the defective mtDNA accumulates in both germline and somatic tissue during post-reproductive life.

### Age-dependent increase in mtDNA*^uaDf5^* burden is transmitted to progeny

As the mtDNA is inherited strictly through the maternal germline, we posited that the age-dependent increase in the fractional abundance of germline mtDNA*^uaDf5^* might be transmitted to progeny animals. To test this hypothesis, we measured the molar ratio of mtDNA*^uaDf5^* in 200-worm populations of L1 larvae derived from day 1 – day 4 adults (Fig. 3C, Fig. 3 – figure supplement 1B). This analysis led to two key observations: 1) the abundance of mtDNA*^uaDf5^* is reduced during transmission between mother and offspring (average decrease ranging from 6% to 13%), presumably as a result of purifying selection, and 2) the abundance of the defective mtDNA in the offspring correlates with the age of the mothers: the progeny of older mothers contain a markedly higher mtDNA*^uaDf5^*:mtDNA^WT^ molar ratio (4.2:1) than that of younger mothers (2.6:1) (Fig. 3C, D). A similar trend was observed for mother-to-offspring transmission in five mutant strains with altered levels of mtDNA*^uaDf5^* (see below): in all cases, progeny contain lower abundance of mtDNA*^uaDf5^* than their mothers, and progeny of younger adults inherit a lower load of mtDNA*^uaDf5^* than progeny of older adults (Fig. 3C). These results reveal that mtDNA quality control occurs between primordial germ cell proliferation in the female germline and L1 hatching, i.e., during oocyte maturation, embryogenesis, or both.

### The lifespan-determining IIS pathway regulates accumulation of mtDNA*^uaDf5^*

We sought to determine whether the age-dependent accumulation of defective mtDNA is controlled by known molecular mechanisms that drive the aging program in *C. elegans*. The most prominent of these regulatory systems is the highly conserved insulin/IGF-1 (insulin-like growth factor-1) pathway (IIS), which performs a pivotal regulatory function in aging and longevity [28,82,83] (Fig. 4 – figure supplement 1). Abrogation of the IIS signaling pathways, for example, as a result of mutations in the gene encoding the IIS receptor (DAF-2, in *C. elegans*), results in marked slowing of the aging program and extension of lifespan in worms, flies, and mice [84–87]. The IIS pathway also functions in a broad set of other processes including, in *C. elegans*, activation of two stages of developmental arrest, or diapause, at the L1 larval stage and in formation of the dispersal form, the dauer larva, as well as in the control of germline proliferation, stress resistance, fat metabolism, and neuronal/behavioral programs [28]. It was also reported that inhibition of the IIS pathway rescues various fitness parameters in a mtDNA mutator strain which contains a faulty mtDNA polymerase [29], consistent with a possible role in mtDNA quality control.

We found that two mutant alleles of *daf-2*, which reduce rates of aging and increase lifespan, result in dramatically decreased mtDNA*^uaDf5^*:mtDNA^WT^ molar ratios from 2.8:1 to as low as 0.3:1 (0.3:1 for *daf-2(e1391*); 0.8:1 for *daf-2(e1370)*; Fig. 4A; see Suppl. Table 2 for a list of lifespan mutants used in the analyses). Thus, the lifespan-extending effects of *daf-2* mutations are strongly correlated with diminished abundance of defective mtDNA, to the extent that it becomes the minor species of mtDNA.

**Figure 4:**
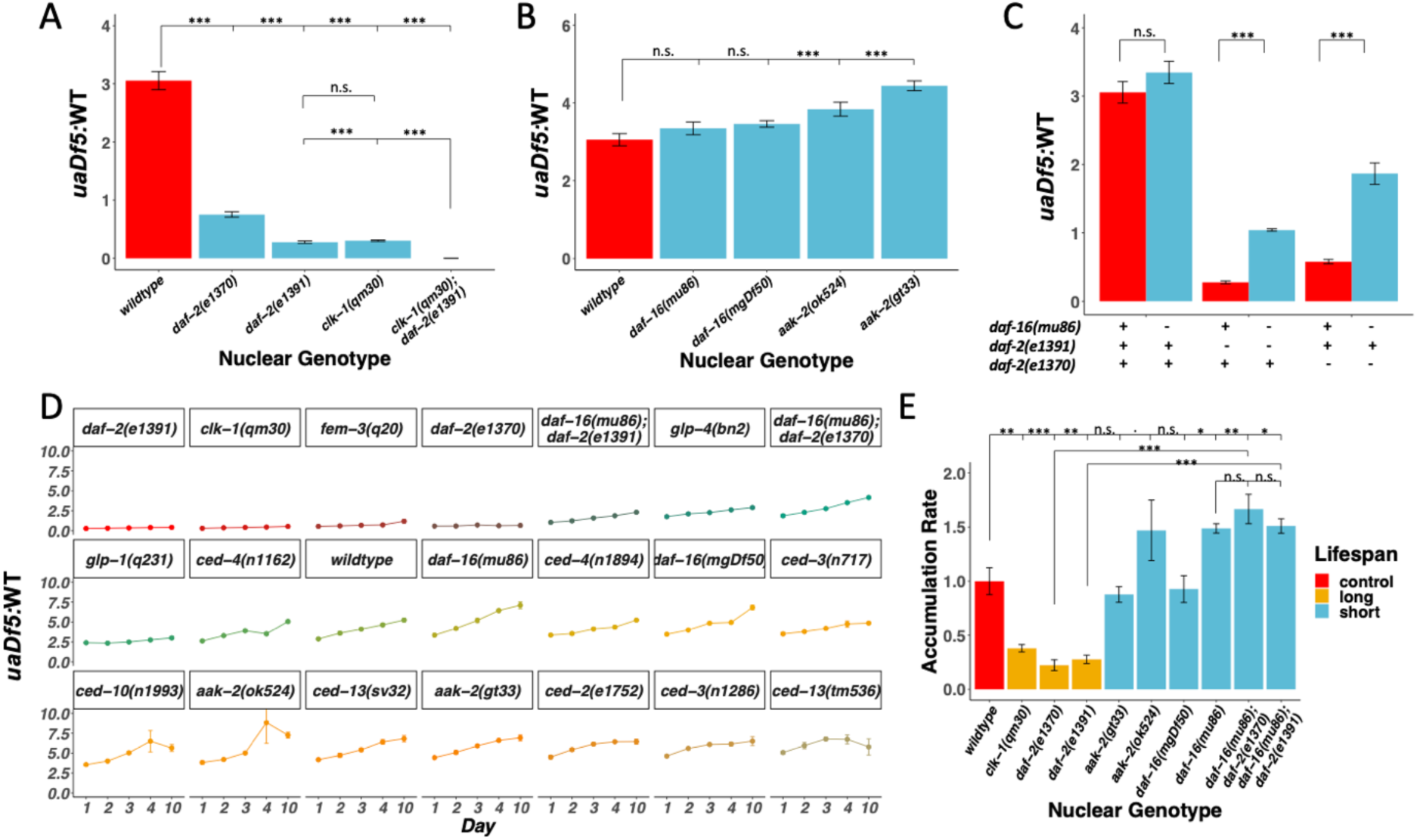
Lifespan mutants have both a lower steady-state level and accumulation rate of mtDNA*^uaDf5^*. **(A-C)** Analysis of molar ratio of mtDNA*^uaDf5^* in day 1 adults of various mutant backgrounds. **(A)** Analysis of steady-state mtDNA*^uaDf5^* levels in long-lived mutants. **(B)** Analysis of steady-state mtDNA*^uaDf5^* levels in short-lived mutants, showing synergistic activity on mtDNA*^uaDf5^* removal capacity in the *daf-2(e1391) clk-1(qm30)* double mutant. **(C)** Analysis of steady-state mtDNA*^uaDf5^* levels in *daf-2(-)* single and *daf-16(-); daf-2(-)* double mutants, showing a partial rescue of *daf-2(-)* phenotype by *daf-16(-)*. **(D)** Analysis of the molar ratio of mtDNA*^uaDf5^* in aging adults in 21 different nuclear backgrounds shows a consistent accumulation trend. **(E)** Summary of the rate of increase for the lifespan regulation mutants, showing that *daf-16* rescues the *daf-2* accumulation rate phenotype. The normalized accumulation rate was calculated by fitting a regression line for each trial and then dividing the slope of the regression line by the slope of the averaged regression line found in a wildtype background. For **all**, n=3 or more replicates of 200 worm populations for each genotype and stage. For **A and E,** statistical analysis was performed using one-way ANOVA with Tukey correction for multiple comparisons. For **C**, statistical analysis was performed using one-way ANOVA. For **B**, statistical analysis was performed using one-way ANOVA with Dunnett’s correction for multiple comparisons. Error bars represent SEM. (*** p < 0.001, ** p < 0.01, * p < 0.05, . p < 0.1, n.s. not significant).

The DAF-2 receptor acts by antagonizing the DAF-16/FoxO transcription factor, the major effector of IIS downstream, in response to insulin-like ligands. Thus, removal of *daf-16* function reverses the lifespan-extending effects of *daf-2(-)* mutants. We tested whether the DAF-2 → DAF-16 pathway similarly functions in mtDNA purifying selection. We found that eliminating DAF-16 in two *daf-16* mutants results in slightly increased, albeit not statistically significant, mtDNA*^uaDf5^*:mtDNA^WT^ molar ratios (3.3:1 for *daf-16(mu86)*; 3.5:1 for *daf-16(mgDf50)*) (Fig. 4B). Further, we found that removal of DAF-16 in the *daf-16(mu86)* mutant suppressed the decreased mtDNA*^uaDf5^*:mtDNA^WT^ molar ratios observed in two *daf-2* mutants, from 0.3:1 for *daf-2(e1391)* to 1:1 for *daf-16(mu86); daf-2(e1391)* and from 0.6:1 for *daf-2(e1370)* to 1.8:1 for *daf-16(mu86); daf-2(e1370)*, consistent with observations reported in a recent study [88]. While the *daf-2(-)* effect on mtDNA*^uaDf5^* levels is largely dependent on DAF-16, neither double mutant restored mtDNA*^uaDf5^* levels to those seen in animals with a fully intact IIS pathway, suggesting that other DAF-2 targets might participate in removal of defective mtDNA (Fig. 4C). We found, conversely, that two mutations that reduce lifespan by eliminating the function of AAK-2 (AMP activated kinase-2), a conserved factor acting in the IIS pathway [28, 89], result in elevated mtDNA*^uaDf5^*:mtDNA^WT^ molar ratios as high as 4.4:1 compared to 2.8:1 in wildtype animals (3.8:1 for *aak-2(ok524)* and 4.4:1 for *aak-2(gt33)*) (Fig. 4B). These findings demonstrate that alterations in the IIS pathway coordinately affect both lifespan and accumulation of defective mtDNA and that the DAF-2/DAF-16/AAK-2 axis acts similarly in both processes.

### Synergistic effect of multiple aging pathways on mtDNA*^uaDf5^* accumulation

In addition to IIS, other molecular regulatory pathways independently contribute to the rate of aging. These include CLK-1, a mitochondrial hydroxylase that functions in the pathway for ubiquinone synthesis [90, 91]. *clk-1* mutants with a wildtype mitochondrial genome have been shown to contain levels of mtDNA that are elevated by 30%, perhaps as the result of a compensatory process that increases demand on mitochondrial abundance, or the action of CLK-1 as a regulator of mtDNA abundance in response to energy availability within the cell [92]. As with long-lived *daf-2* mutants, we found that long-lived *clk-1(qm30)* mutants showed a greatly diminished mtDNA*^uaDf5^*:mtDNA^WT^ molar ratio of 0.3:1 (Fig. 4A), comparable to that in *daf-2* mutants; again, mtDNA*^uaDf5^* is the minor species in these animals. As the IIS pathway and CLK-1 appear to act separately in controlling lifespan, we postulated that elimination of both mechanisms might further reduce levels of the defective mtDNA. Indeed, we found that mtDNA*^uaDf5^* was completely eliminated in *daf-2(e1391) clk-1(qm30)* double mutants, revealing a strongly synergistic effect between the two age-determining systems (Fig. 4A). Thus, distinct regulatory pathways for longevity modulate the abundance of mtDNA*^uaDf5^* by apparently different mechanisms and elimination of the two pathways abrogates its maintenance.

### Age-dependent accumulation rate of mtDNA*^uaDf5^* strongly correlates with genetically altered rates of aging

Our findings that the steady-state abundance of mtDNA*^uaDf5^* increases with maternal age and that mutants with increased lifespan show lower levels of the defective mtDNA raised the possibility that purifying selection is subject to the same control as aging clocks. To assess this potential connection, we analyzed the time-dependent rates of mtDNA*^uaDf5^* accumulation in animals with genetic backgrounds that alter the aging clock. Analysis of 21 different genetic backgrounds over the first ten days of adulthood revealed that the age-dependent progressive accumulation of mtDNA*^uaDf5^* is a consistent phenomenon (Fig. 4D). Comparison of long-lived mutants and wildtype using a linear regression model revealed a striking positive correlation between aging rate and age-dependent rate of accumulation of mtDNA*^uaDf5^*: all long-lived mutants in either the IIS pathway or *clk-1* accumulate mtDNA*^uaDf5^* at a substantially slower rate as they age than do wildtype animals (Fig. 4E, Fig. 4 – figure supplement 2A, B). Conversely, we analyzed six short-lived IIS pathway mutant combinations and found that the *aak-2(ok524)* and *daf-16(mu86)* single mutants and *daf-16(-);daf-2(-)* double mutants all showed increased rates of mtDNA*^uaDf5^* accumulation (Fig. 4E, Fig. 4 – figure supplement 2C, D). These observations suggest that in both slower-aging and faster-aging strains, the rate of accumulation of deleterious mtDNA is a predictor of aging rate. In the two exceptional cases, the *aak-2(gt33)* and *daf-16(mgDf50)* single mutants, we did not observe an increased accumulation rate compared to wildtype; however, the mtDNA*^uaDf5^* levels are consistently higher than in these two mutants than in wildtype at all stages (Fig. 4E, Fig. 4 – figure supplement 2D) and thus diminished removal of mtDNA*^uaDf5^* overall correlates with decreased lifespan in these mutants as well. The greater mtDNA*^uaDf5^* accumulation rates seen in the short-lived animals is not attributable to the higher steady-state levels *per se*, as the rates of accumulation of defective mtDNA observed in the PCD mutants with higher mtDNA*^uaDf5^* levels show no correlation with the steady-state levels (Fig. 4 – figure supplement 2 E-H); rather these increased rates appear specifically to be a property of the shortened lifespan mutants.

Consistent with a relationship between aging rates and accumulation of defective mtDNA, we found that brood size is decreased and embryonic lethality is increased in short-lived *daf-16(-)* mutants compared to those in a wildtype nuclear background, while *uaDf5* does not impact either fitness parameter in the long-lived *clk-1* mutant (Fig. 4 – figure supplement 3). These results are consistent with the possibility that longevity pathways modulate fitness in part by regulating mitochondrial homeostasis.

### Evidence for late adulthood-specific mechanisms for removal of mtDNA*^uaDf5^*

We obtained evidence that defective mtDNA is more effectively removed in offspring of aging adults that carry an unusually high burden of mtDNA*^uaDf5^*. The offspring of day 4 adults in those strains (“high” strains) with significantly higher steady-state fractional abundance of mtDNA*^uaDf5^* showed significantly greater rates of reduction of the defective mtDNA (reduction from 5.4:1 to 3.3:1 in *atfs-1(et15)*, from an extraordinarily high 6.5:1 to 2.8:1 in *ced-10(n1993),* and from 6.4:1 to 3.3:1 in *ced-13(sv32)*) compared to offspring of (“low” strain) mothers with lower steady-state fractional abundance of the mutant mtDNA (reduction from 4.7:1 to 4.2:1 in WT, from 3.9:1 to 2.8:1 in *ced-3(n2454),* and from 3.2:1 to 2.5:1 in *ced-4(n1162)*). Remarkably, therefore, day 4 progeny from “high” strains actually inherit a *lower* mtDNA*^uaDf5^* load than their siblings born from day 1-3 mothers (Fig. 3C, 5A). Indeed, we found a strong correlation (r^2^=0.61, p < 0.001) between the steady-state level of mtDNA*^uaDf5^* in mothers and the capacity for its removal between mother and progeny during day 4 of adulthood (Fig. 5B). These results raise the possibility that very high levels of mtDNA*^uaDf5^* in older mothers activate additional mtDNA purifying selection independent of the UPR^MT^ and PCD machinery, thereby ensuring that progeny are not overloaded with defective mitochondria.

**Figure 5:**
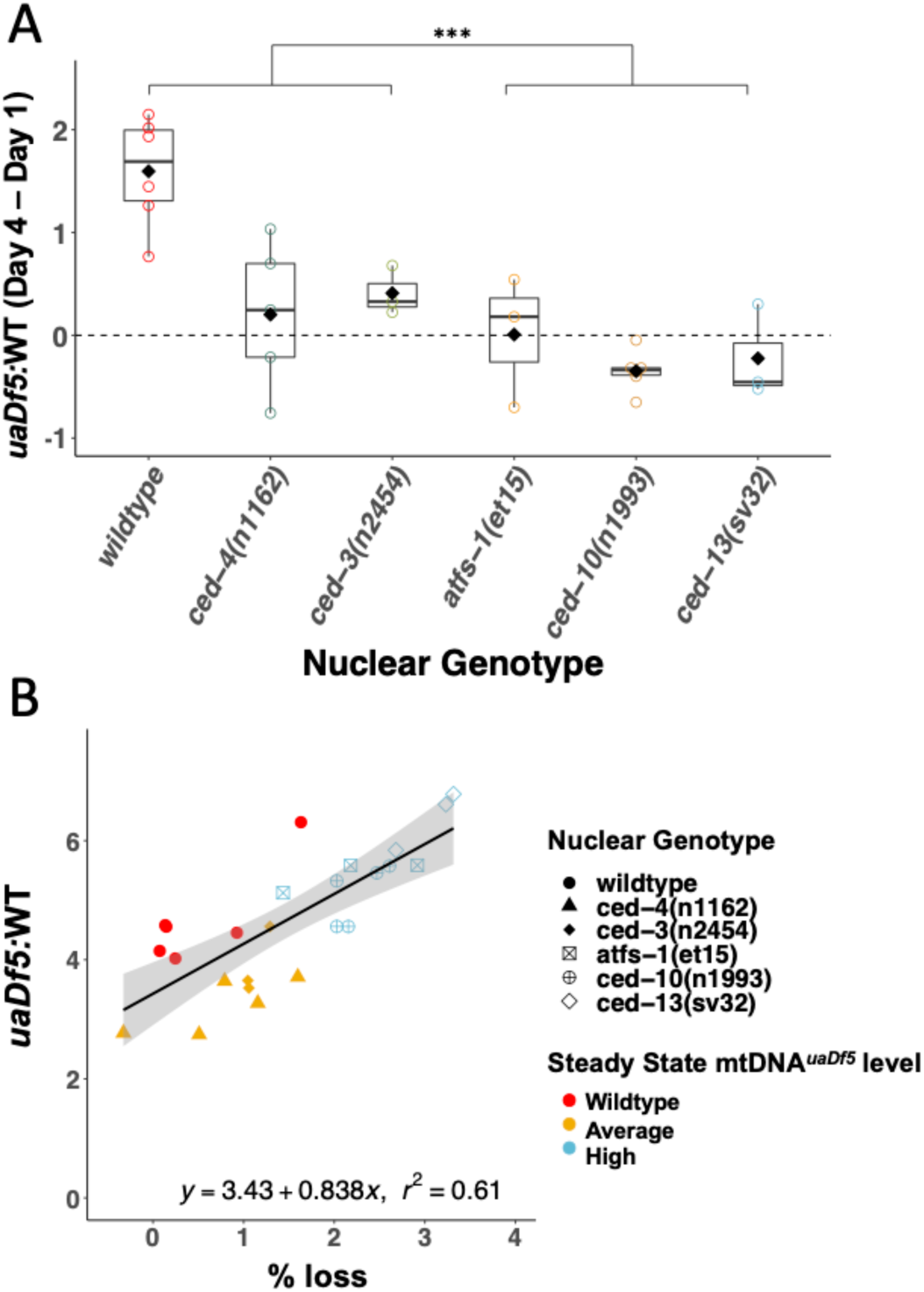
Evidence for late adulthood-specific mechanisms for removal of mtDNA*^uaDf5^*. **(A)** Subtracting *uaDf5:*WT in progeny from day 1 adults from progeny of day 4 adults shows that day 4 F1-L1s tend to have higher mtDNA*^uaDf5^* burden than their day 1 siblings, but this is no longer the case in nuclear backgrounds that result in a significantly higher steady-state levels of mtDNA*^uaDf5^* in the adult. n=3 or more replicates for each genotype and statistical analysis was performed using the Mann-Whitney test. **(B)** Comparison of the molar ratio of mtDNA*^uaDf5^* in day 4 adult mothers to the absolute % removal of mtDNA*^uaDf5^* from mother to offspring shows a positive correlation. (*** p < 0.001).

## Discussion

We have obtained several lines of evidence indicating that regulators of PCD and the aging program function in mtDNA quality control and accumulation of defective mtDNA in *C. elegans*. We report eight major findings: 1) regulators of germline PCD are required for effective removal of deleterious mtDNA from the germline; 2) the cell death machinery functions in either a non-canonical cell death pathway or in a non-apoptotic role to mediate mitochondrial purifying selection; 3) the CSP-1 caspase has as strong of an effect on mitochondrial purifying selection as the major PCD regulator CED-3; 4) mtDNA*^uaDf5^* progressively accumulates in the germline as adults age; 5) this age-dependent accumulation of mtDNA*^uaDf5^* is transmitted to progeny; however, the burden of the defective mtDNA is lower in offspring than mothers suggesting intergenerational purifying selection; 6) two separate aging pathways, the IIS and CLK-1 pathways, act synergistically to regulate mtDNA*^uaDf5^* levels; longer-lived mutants show reduced levels of the defective mtDNA while shorter-lived mutants show increased levels compared to otherwise wildtype animals; 7) the rate of mtDNA*^uaDf5^* accumulation is inversely correlated with lifespan in aging mutants; 8) intergenerational removal of mtDNA*^uaDf5^* occurs more effectively during transmission from older mothers with high burden of the defective mtDNA.

Previous reports demonstrated that UPR^MT^ limits *uaDf5* clearance, and that eliminating the UPR^MT^-mediating transcription factor, ATFS-1, lowers mtDNA*^uaDf5^* abundance [25, 27]. Our identification of a second mutation in the *uaDf5* mutant that results in premature truncation of the ND4 gene product raises the possibility that expression of both the truncated ND4 and the ND1-CYTB fusion protein might together activate UPR^MT^. The possibility that it is the production of aberrant polypeptides resulting from these mutations that trigger this response will require analysis of additional mtDNA mutants.

### Mitochondrial deletion mutant *uaDf5* as a model for mitochondrial disease

We found that *uaDf5* affects brood size, embryonic lethality, and developmental rate, highlighting its use as a model for investigating mitochondrial diseases. The reduced brood size in *uaDf5*-bearing animals might reflect diminished germ cell proliferation, as mitochondria have been implicated in progression of germline maturation [93, 94]. Alternatively, the defective mtDNA might trigger hyperactivation of the germline PCD pathway that specifically removes germ cells with the highest burden of defective mtDNA, as suggested by our results, resulting in the survival of fewer mature oocytes. The increased embryonic lethality in the *uaDf5* strain may be a consequence of a genetic bottleneck effect, leading to rapid differences in mtDNA allele frequencies [95–97] and a subpopulation of oocytes containing levels of mtDNA*^uaDf5^* that exceed a threshold required for viability.

We were surprised to find, in contrast to a previous report [98], that mtDNA*^uaDf5^* did not alter lifespan. Given that mitochondrial mutations are often coupled with compensatory mutations in the nuclear genome [99–103], one possible explanation for this discrepancy might be that a compensatory nuclear mutation exists in the strain analyzed, diminishing the impact of the defective mitochondrial genome. We note, however, that we backcrossed the *uaDf5* strain extensively to the laboratory reference strain N2 prior to performing the reported analyses. It is conceivable that although we observed no effect in the lab, *uaDf5* might alter lifespan under natural conditions. Exposure to increased stress from growth in the wild might be less tolerated in animals bearing mtDNA*^uaDf5^*, as has been observed with other mitochondrial mutants [104], resulting in diminished lifespan.

### Caspases and cell death machinery regulate mitochondrial purifying selection

Our results lend support to the hypothesis that germline PCD mechanisms may be used to cull germline cells with defective mtDNA (Fig. 6A). Developmentally programmed cell death and cell death in response to genotoxic stress are mediated by caspases upon activation by the *C. elegans* octameric apoptosome which is formed when the inhibition of CED-4 by CED-9 is disrupted through binding of BH-3-only proteins EGL-1 and CED-13 in the soma and germline, respectively. We propose that in response to mitochondrial genotoxic stress (increased mtDNA*^uaDf5^* load), caspases CED-3 and CSP-1 are activated by the BH-3 only domain protein CED-13 thereby triggering mitochondrial purifying selection independent of CED-9 and the CED-4 apoptosome, either through germ cell PCD or a non-apoptotic role of these caspases (Fig. 6A).

**Figure 6:**
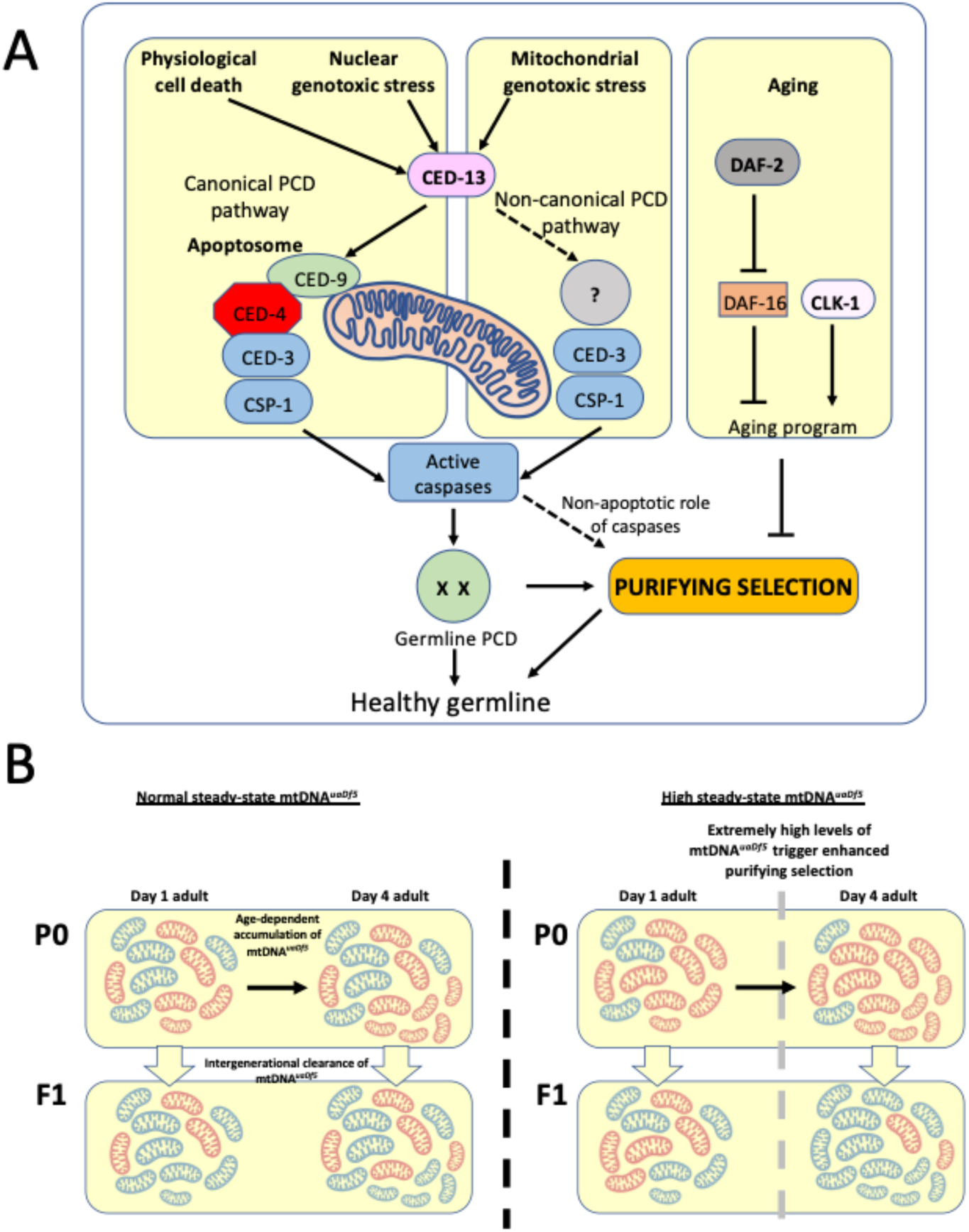
Regulation of mtDNA*^uaDf5^* accumulation and transmission by the PCD and aging pathways. **(A)** Our results suggest that CED-3 and CSP-1, which are activated by BH-3 only protein CED-13, function cooperatively to promote mitochondrial purifying selection, independent of CED-9 and the CED-4 apoptosome. The clearance of mtDNA*^uaDf5^* may therefore involve induction of a non-canonical germline PCD mechanism or non-apoptotic action of the CED-13/caspase axis. Additionally, IIS and CLK-1 aging pathways act synergistically to regulate mitochondrial purifying selection**. (B)** mtDNA*^uaDf5^* (red) accumulates in the germline relative to mtDNA^WT^ (blue) as adults age and the increased mtDNA*^uaDf5^* levels are transmitted to the progeny, although the mtDNA*^uaDf5^* burden is consistently lower in progeny than mothers. This intergenerational purifying selection is enhanced in the older mothers of mutants with high steady-state mtDNA*^uaDf5^* (e.g., *ced-13*, *ced-10,* and *atfs-1*), suggesting a threshold beyond which a germline PCD-independent mtDNA quality control process may be initiated or enhanced in these older mothers.

Mutations that eliminate the function of caspases that act in PCD result in elevated mtDNA*^uaDf5^* levels. Two mutations affecting the p15 domain of the CED-3 caspase result in a significantly increased molar ratio of *uaDf5*, highlighting the importance of the p15 domain in mtDNA quality control. It is noteworthy that the two mutations that result in a more substantial effect on mtDNA*^uaDf5^* abundance affect CED-3 structure more dramatically: *ced-3(n717)* results in a splicing error, and *ced-3(n1286)* is a nonsense mutation [54]. In contrast, *ced-3(n2454)*, which results in a subtle (statistically insignificant) increase in mtDNA*^uaDf5^,* is a substitution predicted to impart a much weaker effect on the protein structure [54]. The effect of the *ced-3(n718)* allele, which resides in the prodomain of CED-3, suggests that that portion of the protein may act to inhibit mtDNA quality control. Taken together, our results suggest that CED-3 may carry out a specialized activity in mtDNA-activated PCD. We found that a second caspase, CSP-1, which plays a minor role in PCD [56, 57], is also required in mtDNA quality control: a *csp-1(-)* knockout mutation results in increased abundance of mtDNA*^uaDf5^* and enhances the effect of *ced-3(n717)*, suggesting that CSP-1 may play a larger role in removal of defective mtDNA than it does in other forms of PCD. An exciting possibility is that caspases act in mtDNA quality control via a mechanism that is distinct from their normal action in PCD. Such putative roles for caspases and other mitochondrial factors in non-apoptotic mitochondrial quality control may have been co-opted during metazoan evolution with the innovation of PCD.

Analysis of additional PCD components further implicate a role for germline PCD in mitochondrial purifying selection. These include CED-13, the germline-specific PCD effector, and components of the cell corpse engulfment pathway, which activate PCD likely through a complex feedback mechanism that ensures cells destined to die proceed irreversibly through the process [66, 67]. In all cases, mutations in these components result in increased mtDNA*^uaDf5^* levels. In vertebrates, mitochondrial reactive oxygen species (mtROS) trigger apoptosis via the intrinsic mitochondrial pathway [120]. Interestingly, elevated mtROS promotes longevity that is in part dependent on the core cell death machinery in *C. elegans*, involving CED-9, CED-4, CED-3 and CED-13, but not EGL-1 [105]. In that study, the authors reported that the protective effect of the cell death machinery on lifespan was independent of PCD in the soma; however, germline cell death was not characterized. Given that CED-13, and not EGL-1, is the predominant BH3-domain protein functioning in the germline [58] and that ablation of the germline leads to extended lifespan [106], our findings support the possibility that germline progenitors carrying defective mitochondria selectively undergo PCD, ensuring homeostatic mtDNA copy number and health of progeny.

A striking exception to our findings was seen with mutations that eliminate the function of the pro-apoptotic regulator CED-4. Neither the *ced-3(n718)* mutation that disrupts the CED-3 CARD domain, which is involved in recruitment to the apoptosome by stabilizing its interaction with CED-4 [44, 107], nor two *ced-4(-)* mutations, result in increased accumulation of mtDNA*^uaDf5^*, suggesting non-canonical, CED-4-independent activation of CED-3 in mitochondrial purifying selection. It is possible that the CARD mutation (G65R) in the *ced-3(n718)* mutant [54] reduces the fraction of CED-3 in complex with the apoptosome, which might release more of the protein for its role in mitochondrial purifying selection. Interestingly, the CARD linker domain has been found to have an inhibitory effect on the pro-caspase-9 zymogen [108]. *ced-3(n718)* could be effectively acting as a gain-of-function allele in the process of purifying selection, reflected by the lower levels of mtDNA*^uaDf5^* in this mutant background.

It is noteworthy that CED-4 and its mammalian Apaf1 relatives regulate a variety of cellular functions that are unrelated to their activities in PCD. These include cell growth control influenced by DNA damage, centrosomal function and morphology, neuronal regeneration, and inhibition of viral replication [109–112]. In addition, as a result of differential RNA splicing, *ced-4* encodes proteins with opposing activities, generating both an activator and a repressor of apoptosis [113], which further complicates analysis of its action. Thus, it is conceivable that CED-4 might exert opposing effects on purifying selection, reflecting its pleiotropic activities in development and confounding an unambiguous interpretation of its action in this process.

### IIS and CLK-1 synergistically regulate germline accumulation of mtDNA*^uaDf5^* as adults age

We found that most of the increase in the fractional abundance of mtDNA*^uaDf5^* as worms age throughout the period of self-fertility (days 1-4) occurs in the germline. However, the relative amount of the defective mtDNA continues to increase in older animals when the germline is no longer actively proliferating [76], suggesting that mtDNA proliferation also occurs in somatic tissues throughout the aging process. This behavior mirrors the dynamics of mutant mtDNAs observed in other organisms, including human, mouse, rat, and rhesus monkey [35,73,79]. While removal of the germline in worms results in extended lifespan [114, 115], it is not clear whether, or to what extent, this increased lifespan might be attributable to accumulation of mutant mtDNA, since the lack of germline leads to a variety of cellular responses [116, 117], any of which might lead to lifespan extension.

These studies do not reveal whether, or how, aging and accumulation of defective mtDNA are causally linked. However, our findings that steady-state levels of mtDNA*^uaDf5^* are lowered in long-lived mutants (*daf-2* (IIS pathway; [28]) and *clk-1* (mitochondrial function; [118, 119])) and that rates of its accumulation are strongly inversely correlated with lifespan extension through independent pathways, suggests that mtDNA purifying selection mechanisms are influenced by aging programs (Fig. 6A). Further bolstering this potential link is our finding that short-lived mutants (*daf-16* and *aak-2,* both involved in the IIS pathway [28]), show higher steady-state levels of mtDNA*^uaDf5^.* That this effect is greater in *aak-2* mutants than in *daf-16* mutants suggests that the AAK-2 branch of the IIS pathway influences mtDNA quality control more significantly than does the DAF-16 branch. One of the substrates of AAK-2 is SKN-1/Nrf2, a multifaceted transcription factor with roles in stress response and longevity, and one of its isoforms, SKN-1a, localizes to the mitochondrial surface [120], raising the possibility that AAK-2 might influence clearance of defective mtDNA through SKN-1 action. It will be of interest to assess how defective mtDNA might coordinately trigger quality control and stress-response pathways.

Our observation that IIS pathway components and CLK-1 act synergistically on the mtDNA quality control machinery raises the possibility that these two distinct lifespan-regulating pathways converge on a common system for removal of defective mtDNA, such as a global mitochondrial stress response pathway. Candidates for mediating this removal process include mitochondrial fission/fusion [17,20,121], mitophagy [18,19,122], and the UPR^MT^ [22–27], all of which are known to act in mtDNA quality control, as well as modulation of the regulatory pathway for PCD, as suggested by our findings.

### PCD is uncoupled from aging during intergenerational mitochondrial purifying selection in older mothers

Analysis of newly hatched L1 larvae revealed that the relative load of mtDNA*^uaDf5^* is transmitted from mother to offspring, with evidence for intergenerational purifying selection (Fig. 6B). This finding implies that mtDNA quality control occurs between germline stem cell expansion in the mature female germline and L1 hatching, a developmental period that spans many potential stages at which it might occur, including germline PCD, oocyte maturation, and the entirety of embryogenesis. It is conceivable that this selection process acts at multiple stages throughout this developmental window and that the decreased burden of defective mtDNA in newly hatched L1 larvae reflect the summation of a series of sequentially acting processes that incrementally enrich for healthy mtDNA.

The efficacy of intergenerational removal of mtDNA*^uaDf5^* increases in old mothers, including in strains lacking pro-apoptotic regulators. The clearance is particularly precipitous in strains with a very high burden of defective mtDNA as seen in the absence of CED-13, CED-10, and ATFS-1, suggesting a critical threshold beyond which a germline PCD-independent mtDNA quality control process may be triggered in these older mothers (Fig. 6B). One possible explanation for this observation is that an mtDNA purifying selection mechanism that is typically inhibited by germline PCD might be activated in older mothers. This hypothesized purifying selection mechanism might be triggered by the unique cellular environment associated with aging such as increased organelle or macromolecule damage. Alternatively, the effect might be the result of an age-dependent genetic program.

While our study has uncovered new mechanisms acting in mtDNA purifying selection and its relationship to aging, it is of note that some level of mtDNA*^uaDf5^* is maintained in all but the most extreme conditions we have observed (e.g., in the *daf-2(e1391) clk-1(qm30)* double mutant, in which the defective mtDNA is extirpated). This finding raises the possibility that some degree of heteroplasmy, even with defective mtDNA, is not only tolerated, but may be adaptive by providing a degree of evolutionary plasticity. Cells might purposefully allow for limited heteroplasmy as a way of increasing genetic heterogeneity that might prove evolutionarily advantageous. Such heterogeneity may also be essential to allow mtDNA to co-evolve with changes arising in the nuclear genome. It may be that a dynamic balance between active mtDNA purifying selection, including the mechanisms identified here, and the permissibility of limited heteroplasmy, is modulated according to environmental or physiological demands.

## Materials and Methods

### Culturing of nematodes

Nematode strains were maintained on NGM plates as previously described at either 20°C or 15°C for the temperature-sensitive strains [123]. Supplementary Tables 1, 2 and 3 provide details of all strains used in this study. Strains without a JR designation were either provided by the CGC which is funded by NIH Office of Research Infrastructure Programs (P40 OD010440) or were obtained from the Mitani lab (strains with a FX designation or JR strains containing alleles with a tm designation were generated from Mitani lab strains) [124].

### Population collection by age

Upon retrieval of a stock plate for a given strain, three chunks were taken from the stock plate and placed onto three separate large NGM plates to create three biological replicates (“lines”). Each of these lines was chunked approximately each generation to fresh large NGM plates (every 3 days if maintained at 20°C or 25°C, or every four days if maintained at 15°C, being careful to not let the worms starve between chunks). After four generations of chunks, an egg prep was performed on each line (as described previously; [123]) and left to spin in M9 overnight to synchronize the hatched L1s. The next day, each egg prep was plated onto three large seeded plates at an equal density and the worms were left to grow to day 2 adults (second day of egg laying). The day 2 adult worms were egg prepped for synchronization and left to spin in M9 buffer overnight. The next day, each egg prep was plated onto five large NGM plates at equal density. Once the worms reached day 1 of adulthood (first day of egg-laying), one of the plates was used to collect 200 adult worms by picking into 400 μl of lysis buffer, and the remaining adults on the plate were egg prepped for the collection of hatched L1 larvae in 400 μl lysis buffer the following day. The worms on the four remaining plates were transferred to a 40 μm nylon mesh filter in order to separate the adults from the progeny, and the resulting adults were resuspended in M9 and pipetted onto fresh large NGM plates. This process was repeated for the following three days (Day 2-4 of adulthood). Day 5-10 adults were moved to fresh NGM plates every 2^nd^ day using a 40 μm nylon mesh filter, and the resulting day 10 adults were collected in lysis buffer.

### ddPCR

The worm lysates were incubated at 65°C for 4 hours and then 95°C for 30 minutes to deactivate the proteinase K. Each lysate was diluted; 100-fold for 200 worm adult population lyses, 2-fold for 200 worm L1 population lyses, and 25-fold for individual adult lyses. 2 μl of the diluted lysate was then added to 23 μl of the ddPCR reaction mixture, which contained a primer/probe mixture and the ddPCR probe supermix with no dUTP. The primers used were:

WTF: 5’-GAGGGCCAACTATTGTTAC-3’

WTR: 5’-TGGAACAATATGAACTGGC-3’

UADF5F: 5’-CAACTTTAATTAGCGGTATCG-3’

UADF5R: 5’-TTCTACAGTGCATTGACCTA-3’

The probes used were:

WT: 5’-HEX-TTGCCGTGAGCTATTCTAGTTATTG-Iowa Black® FQ-3’

UADF5: 5’-FAM-CCATCCGTGCTAGAAGACAAAG-Iowa Black® FQ-3’

The ddPCR reactions were put on the BioRad droplet generator and the resulting droplet-containing ddPCR mixtures were run on a BioRad thermocycler with the following cycle parameters, with a ramp rate of 2°C/sec for each step:

1. 95°C for 5 minutes
2. 95°C for 30 seconds
3. 60°C for 2 minutes
4. Repeat steps 2 and 3 40x
5. 4°C for 5 minutes
6. 90°C for 5 minutes

After thermocycling, the ddPCR reaction plate was transferred to the BioRad droplet reader and the Quantasoft software was used to calculate the concentration of mtDNA*^uaDf5^* (FAM positive droplets) and mtDNA^WT^ (HEX positive droplets) in each well.

### Lifespan analysis

Confluent large plates were egg prepped and left to spin in M9 overnight for synchronization. The hatched L1s were plated onto large thick plates and allowed to grow to day 2 adults before being egg prepped a second time and left to spin in M9 overnight. The next morning, referred to as day 1 for lifespan determination, L1s were singled out onto small plates. Once the worms started laying eggs, they were transferred each day to a fresh small plate until egg laying ceased, after which the worms remained on the same plate unless bacterial contamination required transfer to a fresh plate. Worms were considered dead if there was no movement after being lightly prodded with a worm pick. Worms that died due to desiccation on the side of the plate were excluded from analysis.

### Brood size and embryonic lethality analysis

Confluent large plates were egg prepped and left to spin in M9 overnight for synchronization. The hatched L1s were plated onto large thick plates and allowed to grow to day 2 adults before being egg prepped a second time and left to spin in M9 overnight. The next morning, L1s were singled out onto small plates. Once the worms started laying eggs, they were transferred each day to a fresh small plate until egg laying ceased. The day after transfer to a fresh plate, unhatched embryos and hatched larvae on the plate from the previous day were counted. This was done for each of the days of laying and the total of unhatched embryos and hatched larvae from all plates from a single worm were tabulated to determine total brood size. To determine embryonic lethality, the total number of unhatched embryos was divided by the total brood size. Worms that died due to desiccation on the side of the plate were excluded from analysis.

### Developmental time course analysis

Confluent large plates were egg prepped and left to spin in M9 overnight for synchronization. The hatched L1s were plated onto large thick plates and allowed to grow to day 2 adults before being egg prepped a second time and left to spin in M9 overnight. The next morning, L1s were singled out onto small plates. The stage of the worms was assayed every 12 hours for the first 72 hours after plating. For determining the stage at 60 hours, L4 worms were divided up into three subgroups based on morphology: young-L4, mid-L4, and late-L4; otherwise, all other staged worms were not divided up into subgroups. Worms that died due to desiccation on the side of the plate were excluded from analysis.

### Genotyping the *w47* allele

DNA was collected from reference strain (N2) and two *uaDf5*-containing strains, LB138 and JR3630. Mitochondrial DNA was extracted and the DNA libraries were prepared using Nextera Kit and then sequenced using an Illumina NextSeq500. Prior to alignment, reads from fastq files were trimmed using Trimmomatic. Trimmed, pair-end reads (2x150) were then mapped to the *C. elegans* assembly reference sequence WBcel235 using Burrows-Wheeler Aligner (BWA) [125]. Picard Tools (http://broadinstitute.github.io/picard/) was used to mark duplicate reads, and SAMtools [126] was used to merge, index and create pile-up format. VarScan [127] was used to call variants, and only variants with minimum coverage of 100 and a minimum variant frequency call of 0.01 were considered for analysis.

### Statistical Analysis

Summary statistics, ANOVA, Mann-Whitney tests, and linear regression were calculated using R v3.4.1. The details of the statistical tests are reported in the figure legends.

## Supplemental Materials

**Fig. 1 – figure supplement 1:**
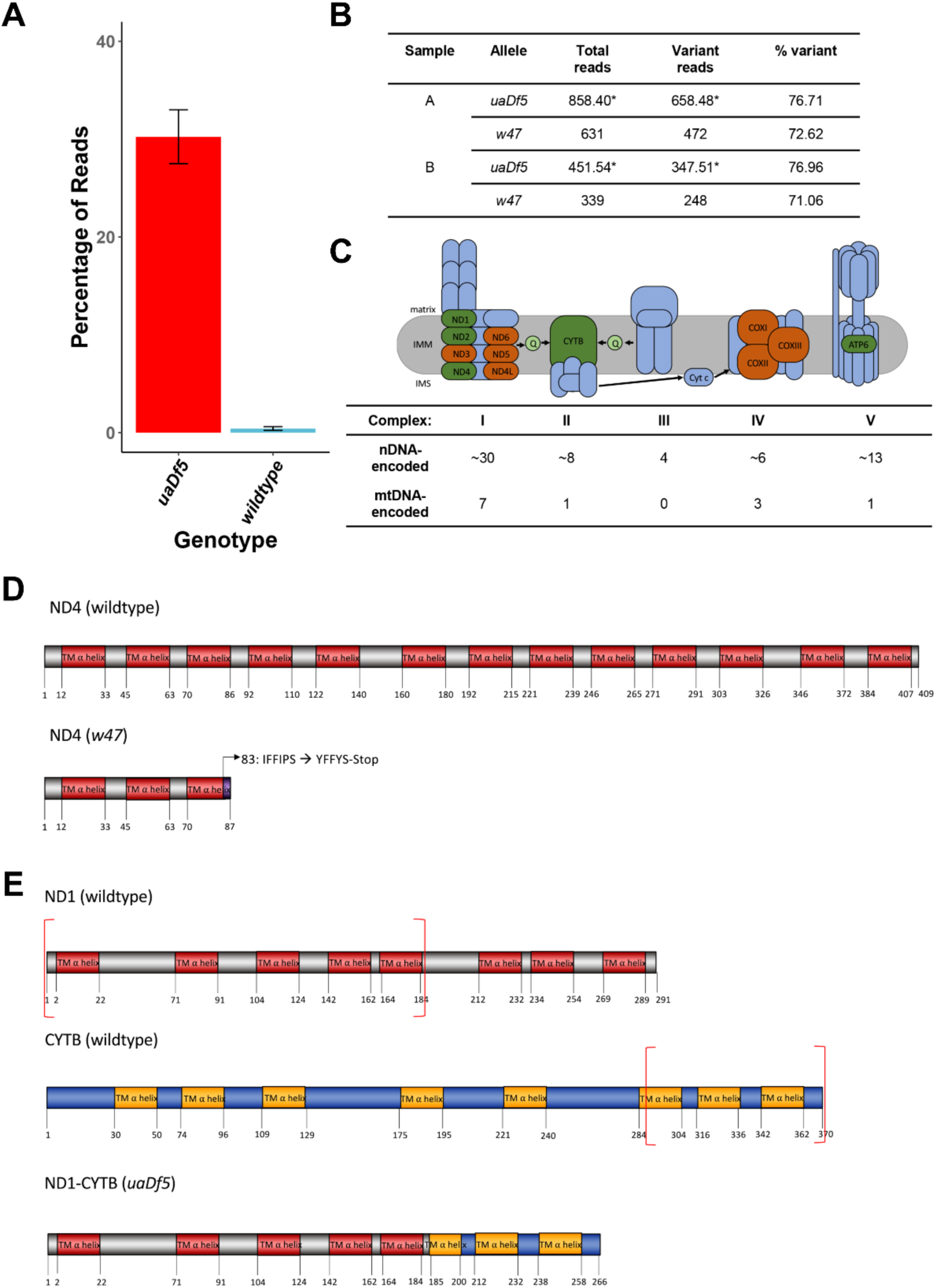
Characterization of the *uaDf5* allele. **(A)** The percentage of reads that mapped to the *w47* insertion in *uaDf5* samples (red) and wildtype samples (blue). Error bars represent SEM. **(B)** Table outlining the percentage of reads that mapped to the *w47* and *uaDf5* mutations in *uaDf5* samples. The number of variant reads was determined by averaging the mapped reads across the deleted region and subtracting that from the average number of reads mapped to the rest of the mtDNA genome (total reads). **(C)** Diagram showing the mitochondrial respiratory chain (MRC) machinery subunits. Blue indicates nDNA-encoded subunits, orange and green indicate mtDNA-encoded subunits. Green indicates those subunits that are knocked out in the *uaDf5* allele (including ND4 which is knocked out by the linked *w47* mutation). **(D)** Diagram showing the likely effect of the *w47* mutation on ND4 protein translation. ND4 is a 409-aa long transmembrane subunit that spans the inner mitochondrial membrane 13 times. The *w47* mutation results in a premature stop codon at position 89, eliminating 10 of the 13 alpha-helix membrane domains. **(E)** Diagram showing the likely effect of the *uaDf5* mutation on ND1 protein translation. ND1 is a 291-aa long transmembrane subunit that spans the inner mitochondrial membrane 8 times and CYTB is a 370-aa long transmembrane subunit that spans the inner mitochondrial membrane 8 times. The *uaDf5* mutation results in a 266 amino acid long fusion protein that connects the first 185 amino acids (and 5 subunits) of ND1 with the last 81 amino acids (and 3 subunits) of CYTB.

**Fig. 1 – figure supplement 2:**
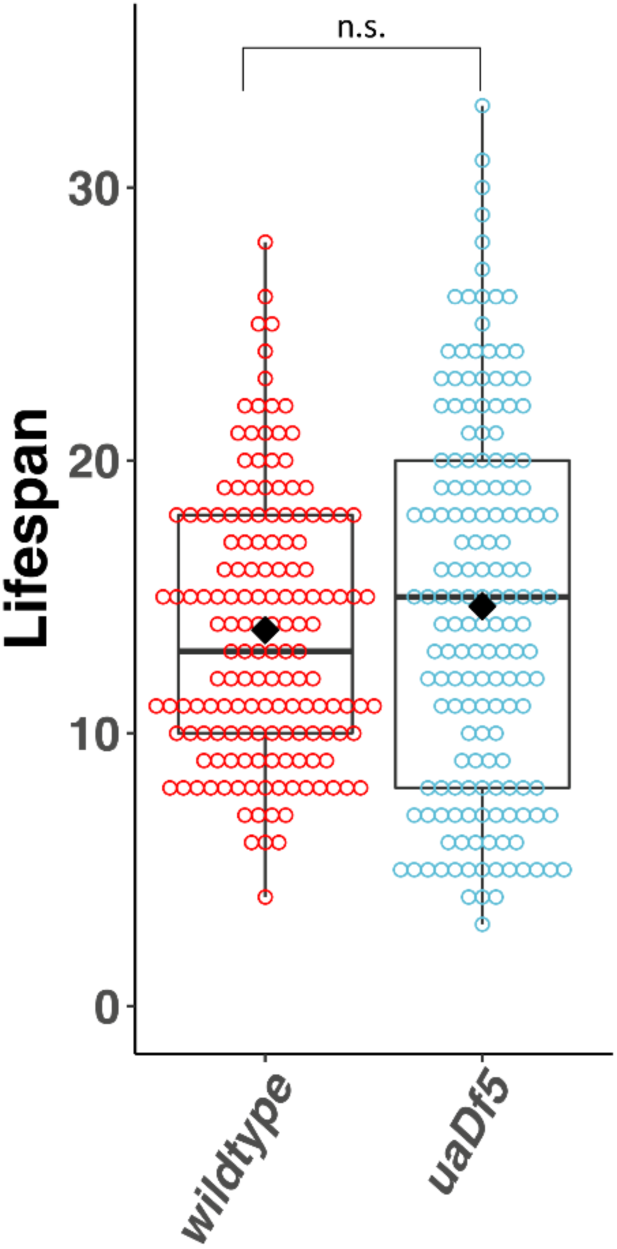
Lifespan analysis of the impact of *uaDf5*. Lifespan analysis of N2 bearing *uaDf5* compared to wildtype N2. Day 1 is defined as the day starved L1s are plated on food. Box plots show median and IQR, and the diamond indicates the mean. Statistical analysis was performed using the Mann-Whitney test. (n.s. not significant).

**Fig. 2 – figure supplement 1:**
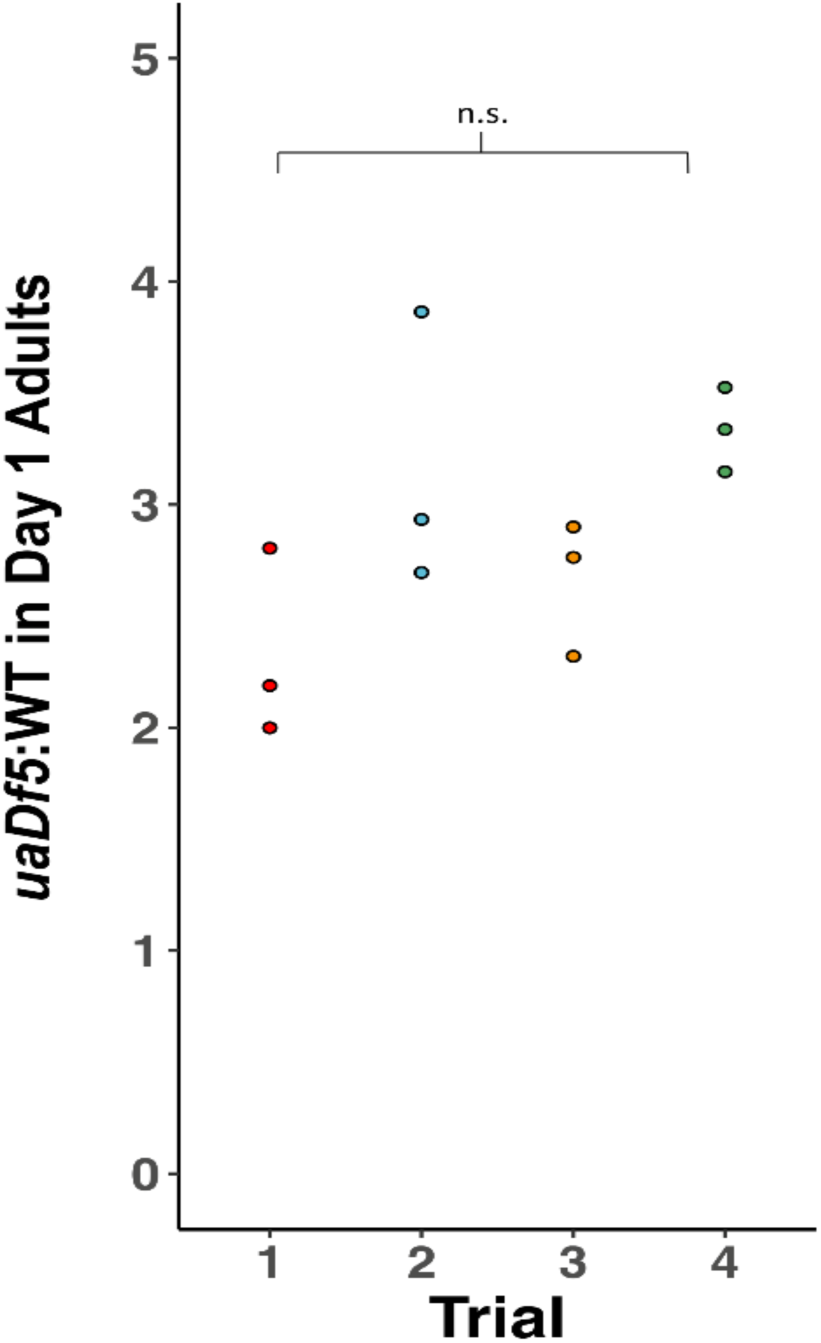
Reproducibility of ddPCR measurement of *uaDf5*. Analysis of the steady-state of *uaDf5* in a wildtype nuclear background shows highly stable steady-state levels. Trials were done months apart on different thaws. Dots represent biological replicates. Statistical analysis was performed using one-way ANOVA with Tukey correction for multiple comparisons. (n.s. not significant).

**Fig. 2 – figure supplement 2:**
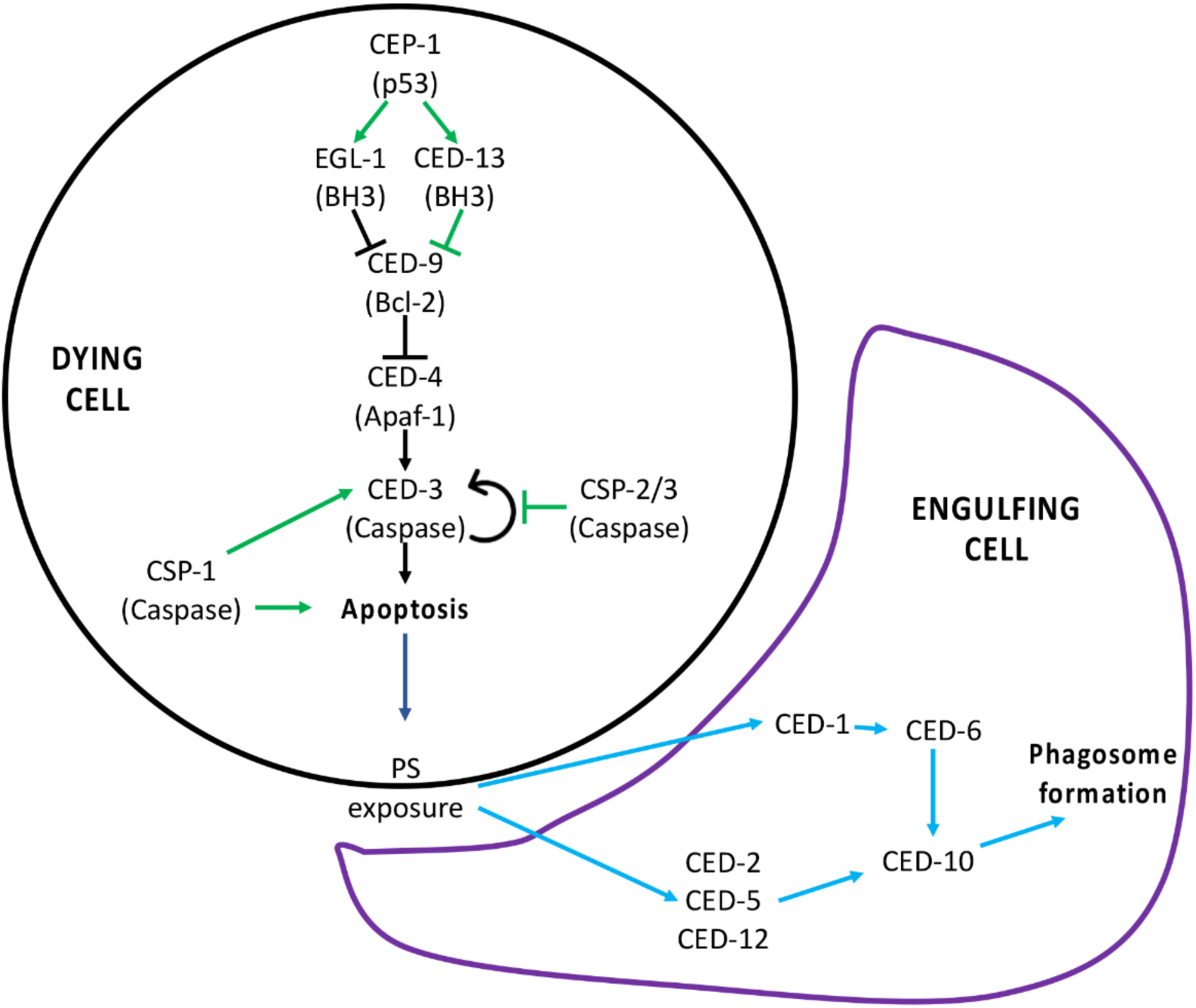
PCD Signaling pathway. **(A)** Signaling pathway for programmed cell death. The canonical pathway is shown with black arrows, the noncanonical pathway is shown with green arrows, and the downstream engulfment pathway is shown with blue arrows. Adapted from [38].

**Fig. 2 – figure supplement 3:**
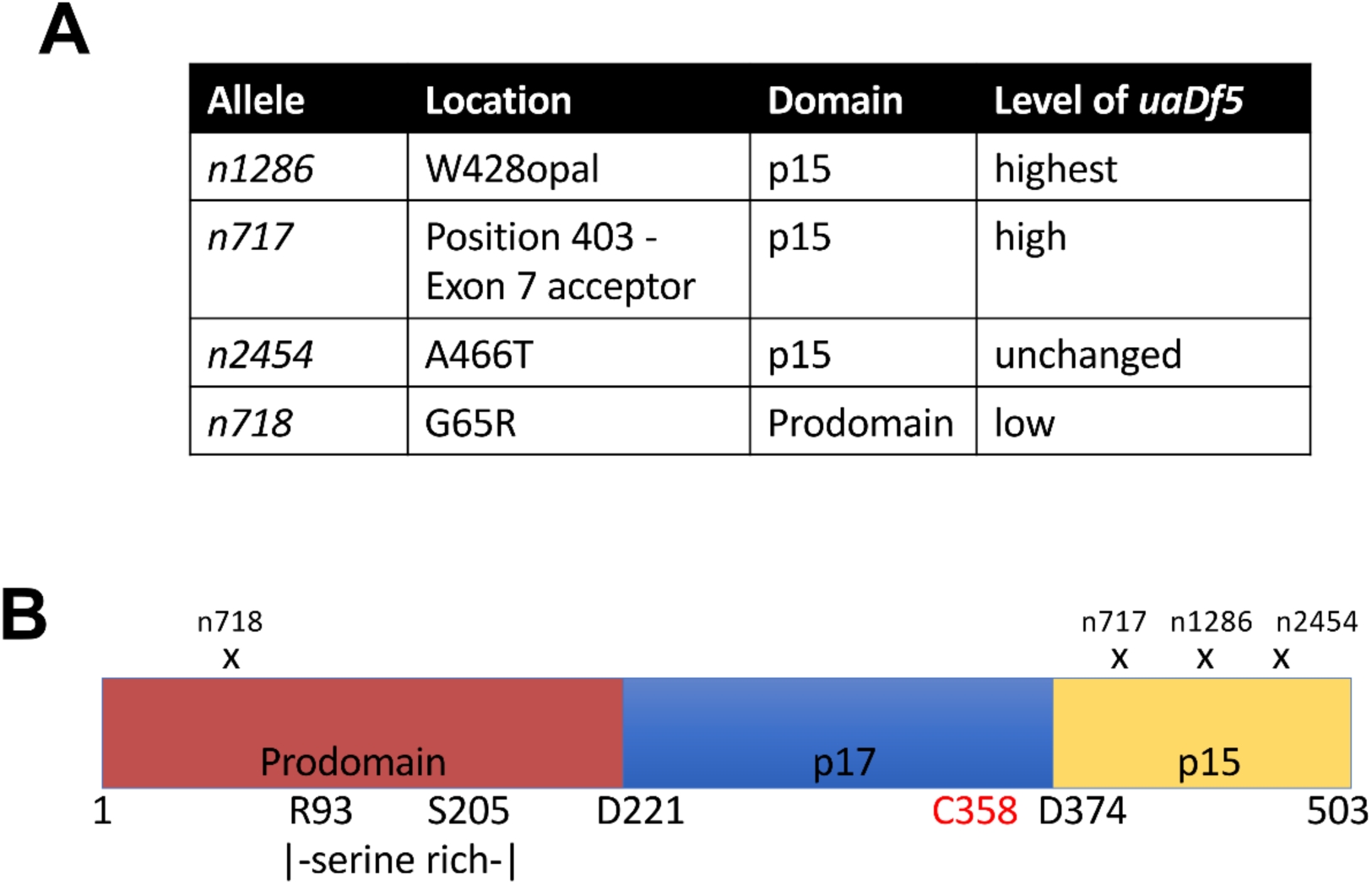
Analysis of the *ced-3* alleles. **(A)** Locations and consequences of the four tested *ced-3* alleles, as well as the measured fractional abundance of *uaDf5*. **(B)** Diagram showing the locations of the mutations for each allele.

**Fig. 2 – figure supplement 4:**
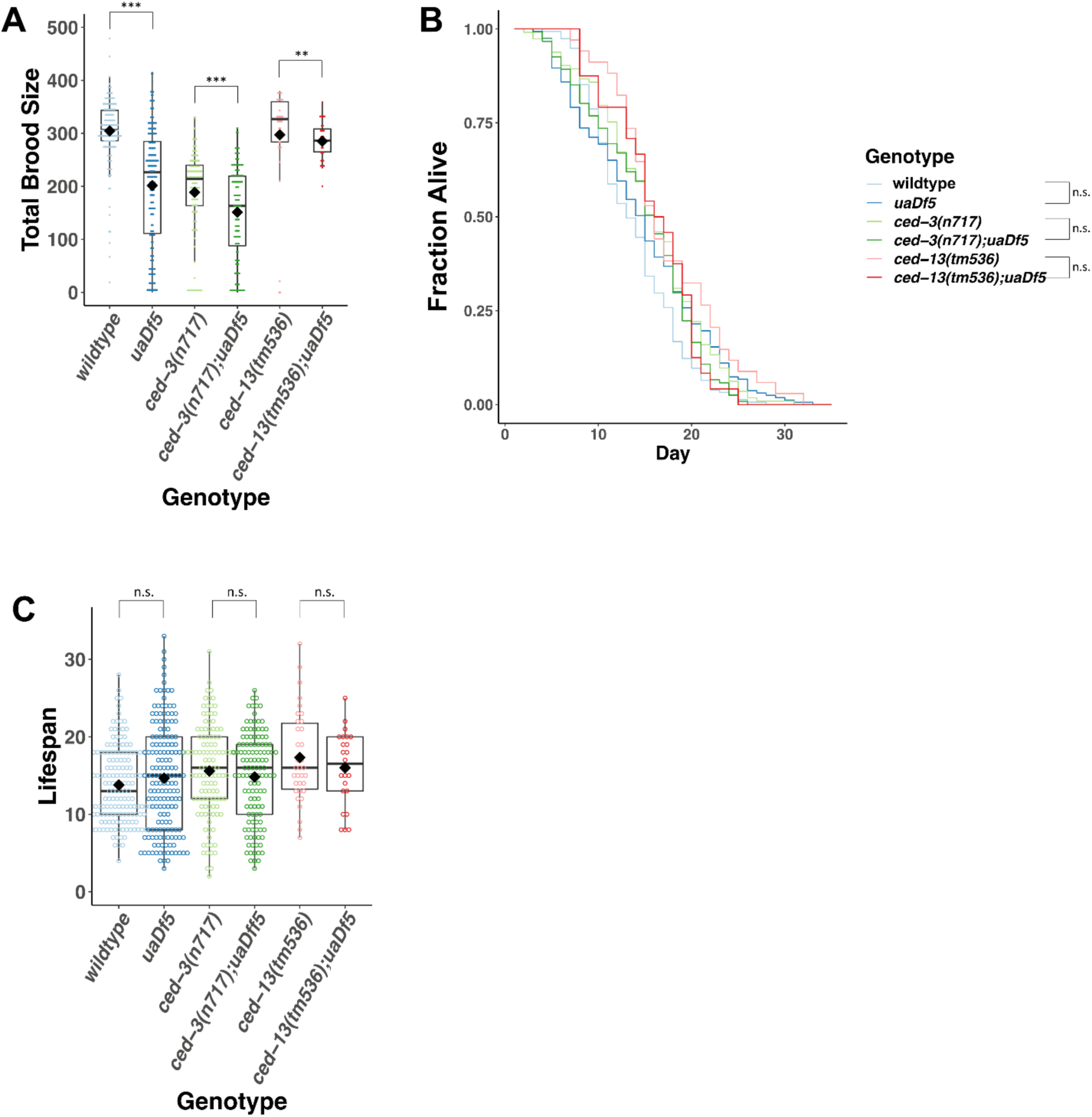
Analysis of the impact of *uaDf5* on fitness parameters in PCD mutants. **(A)** Brood size analysis of PCD mutants. **(B)** Lifespan analysis shows that *uaDf5* does not affect lifespan in both wildtype background and in PCD mutant backgrounds. **(C)** Lifespan analysis shows that *uaDf5* does not affect lifespan in a wildtype nuclear background nor in PCD mutant backgrounds. For **B and C,** day 1 is defined as the day starved L1s are plated on food. For **A and C**, box plots show median and IQR, and the diamond indicates the mean. Statistical analysis was performed using the Mann-Whitney Wilcoxon test. (*** p < 0.001, ** p < 0.01, * p < 0.05, . p < 0.1, n.s. not significant).

**Fig. 3 – figure supplement 1:**
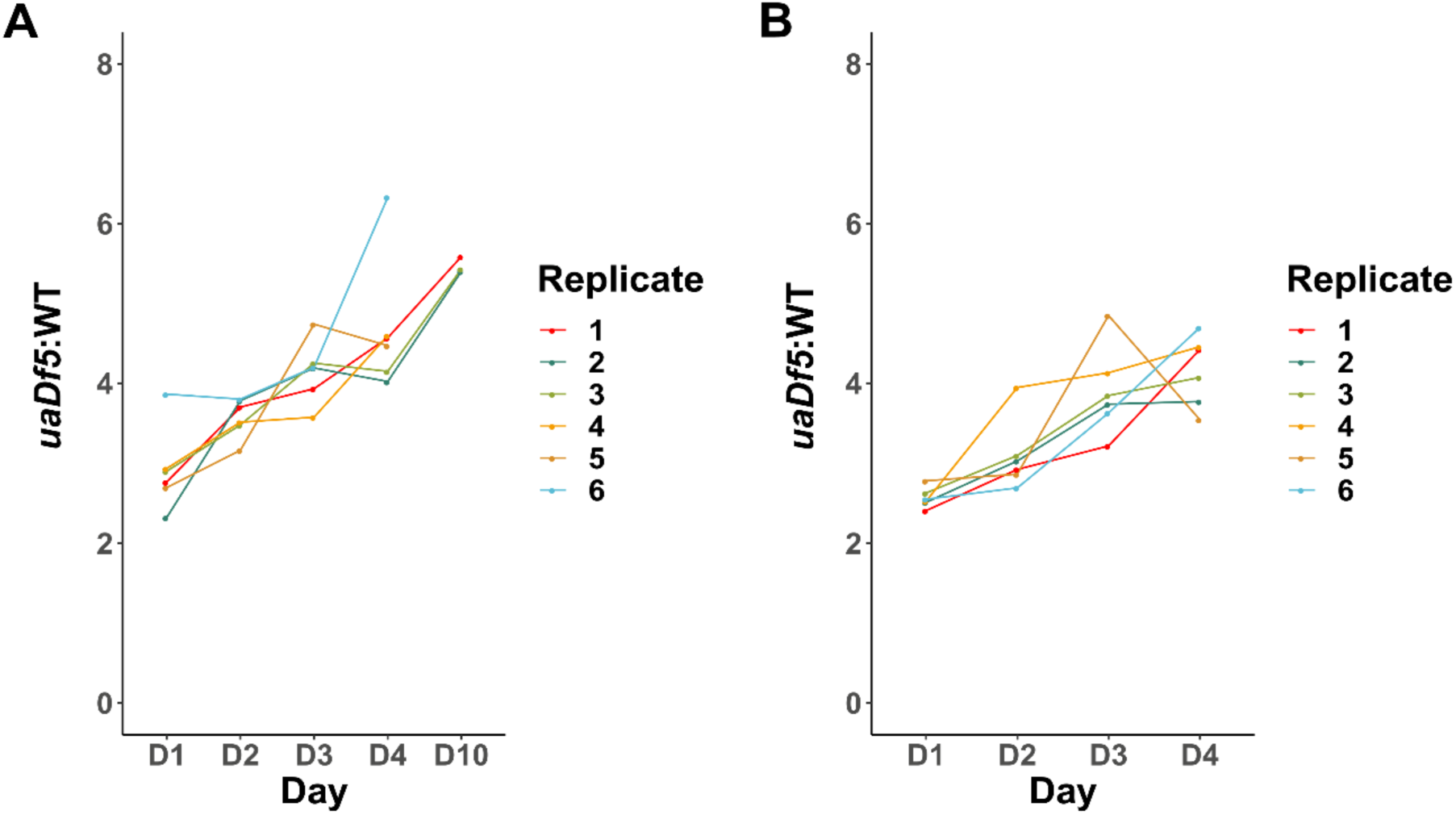
*uaDf5* accumulation in individual lines of adults and progeny. **(A)** Analysis of *uaDf5* accumulation in individual lines of aging P0 adults shows a consistent accumulation trend as adults age. **(B)** Analysis of *uaDf5* accumulation in individual lines of F1-L1 progeny that were born from those mothers shown in panel A show a consistent trend of progeny born from older mothers inheriting a larger *uaDf5* load than their siblings born from younger mothers.

**Fig. 4 – figure supplement 1:**
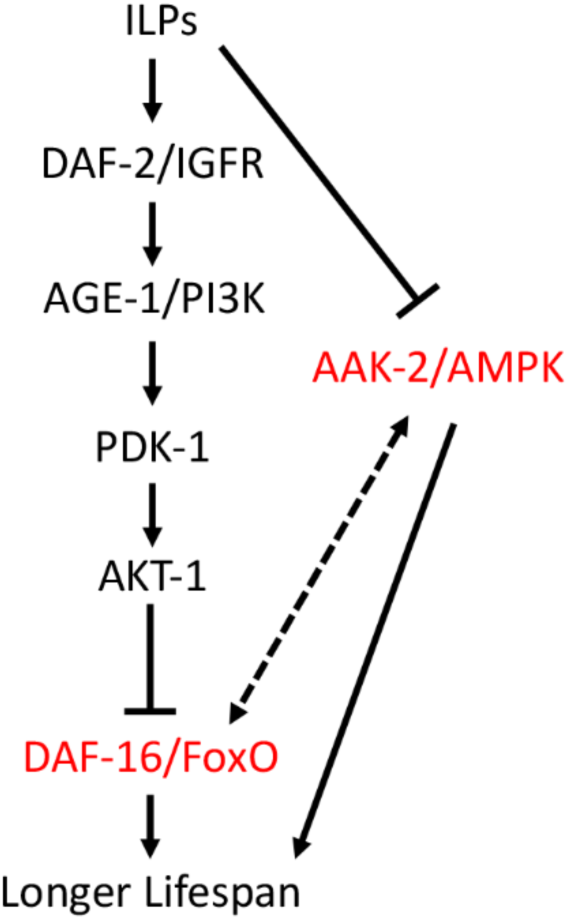
IIS Signaling pathway. Insulin-like peptides (ILPs) bind to DAF-2 and activate the PI3P pathway which prevents nuclear translocation of DAF-16. AAK-2 may phosphorylate and activate DAF-16 transcriptional activity. Loss of *aak-2* or *daf-2* (highlighted in red) reduces lifespan

**Fig. 4 – figure supplement 2:**
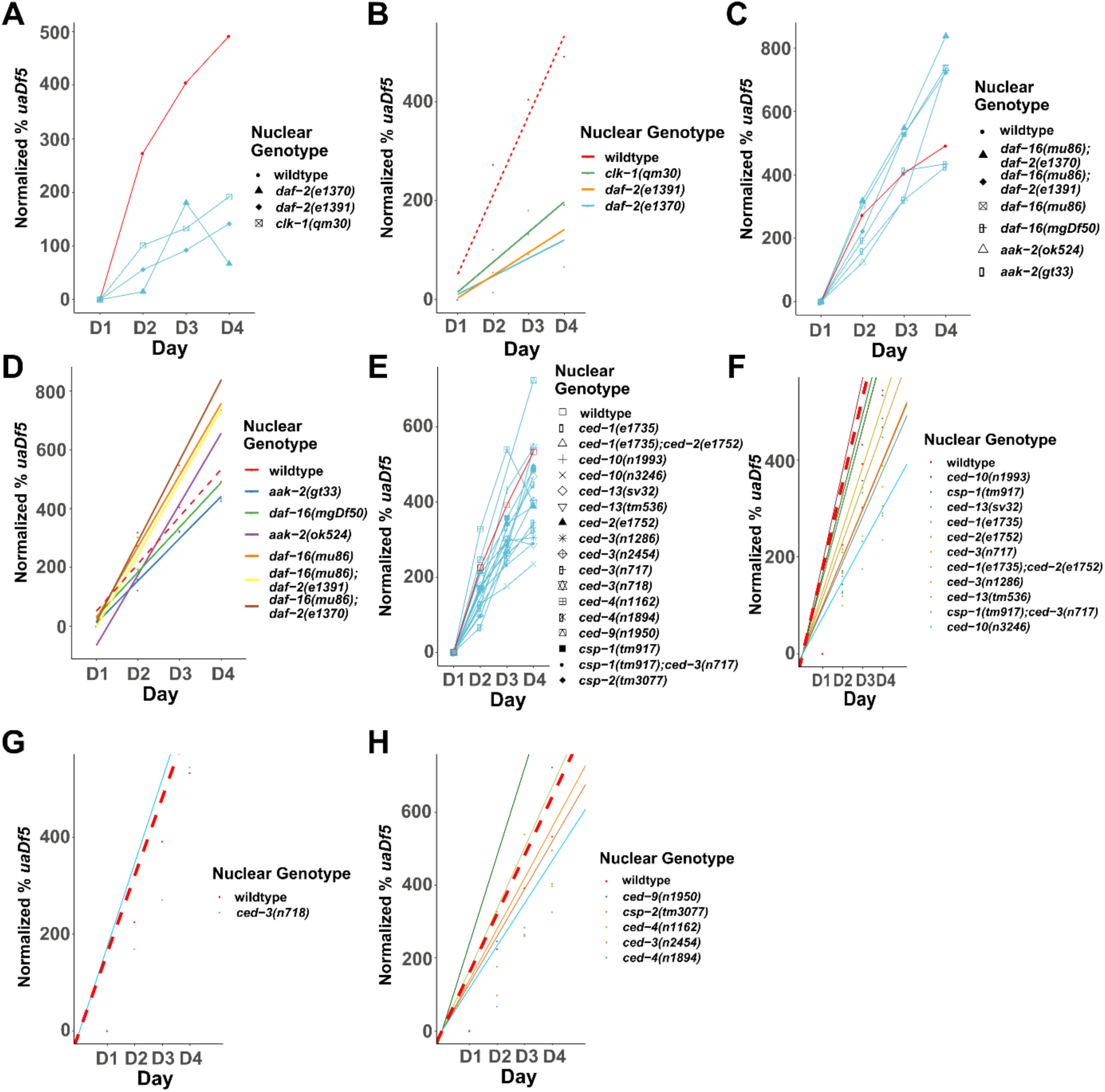
Analysis of the accumulation of mtDNA*^uaDf5^* in PCD and lifespan mutants. Analysis of the normalized fractional abundance *uaDf5* [(% *uaDf5* day x- % *uaDf5* day 1)* % *uaDf5* day 1] in steady-state populations, showing the accumulation rate of mtDNA*^uaDf5^* as worms age from day 1 to day 4 of adulthood. **(A)** Analysis of the accumulation of *uaDf5* in long-lived mutants. **(B)** Analysis of the accumulation of mtDNA*^uaDf5^* in short-lived mutants. **(C-D)** Linear regression analysis of the normalized fractional abundance *uaDf5* [(% *uaDf5* day x- % *uaDf5* day 1)* % *uaDf5* day 1] in steady-state populations, showing the accumulation rate of mtDNA*^uaDf5^* as worms age from day 1 to day 4 of adulthood. **(C)** Analysis of long-lived mutants. **(D)** Analysis of short-lived mutants. **(E)** Analysis of the accumulation of mtDNA*^uaDf5^* in PCD mutants. **(F-H)** Linear regression analysis of the normalized fractional abundance mtDNA*^uaDf5^* [(% *uaDf5* day x- % *uaDf5* day 1)* % *uaDf5* day 1] in steady-state populations, showing the accumulation rate of mtDNA*^uaDf5^* as worms age from day 1 to day 4 of adulthood. **(F)** Analysis of PCD mutants with significantly high steady-state level of mtDNA*^uaDf5^*. **(G)** Analysis of PCD mutants with significantly low steady-state level of mtDNA*^uaDf5^*. **(H)** Analysis of PCD mutants with no significant change in the steady-state level of mtDNA*^uaDf5^*. For **C, D, F, G, and H** dots represent actual datapoints, lines are fitted regression models of the data. For all, n=3 replicates or more of 200 worm populations for each datapoint.

**Fig. 4 – figure supplement 3:**
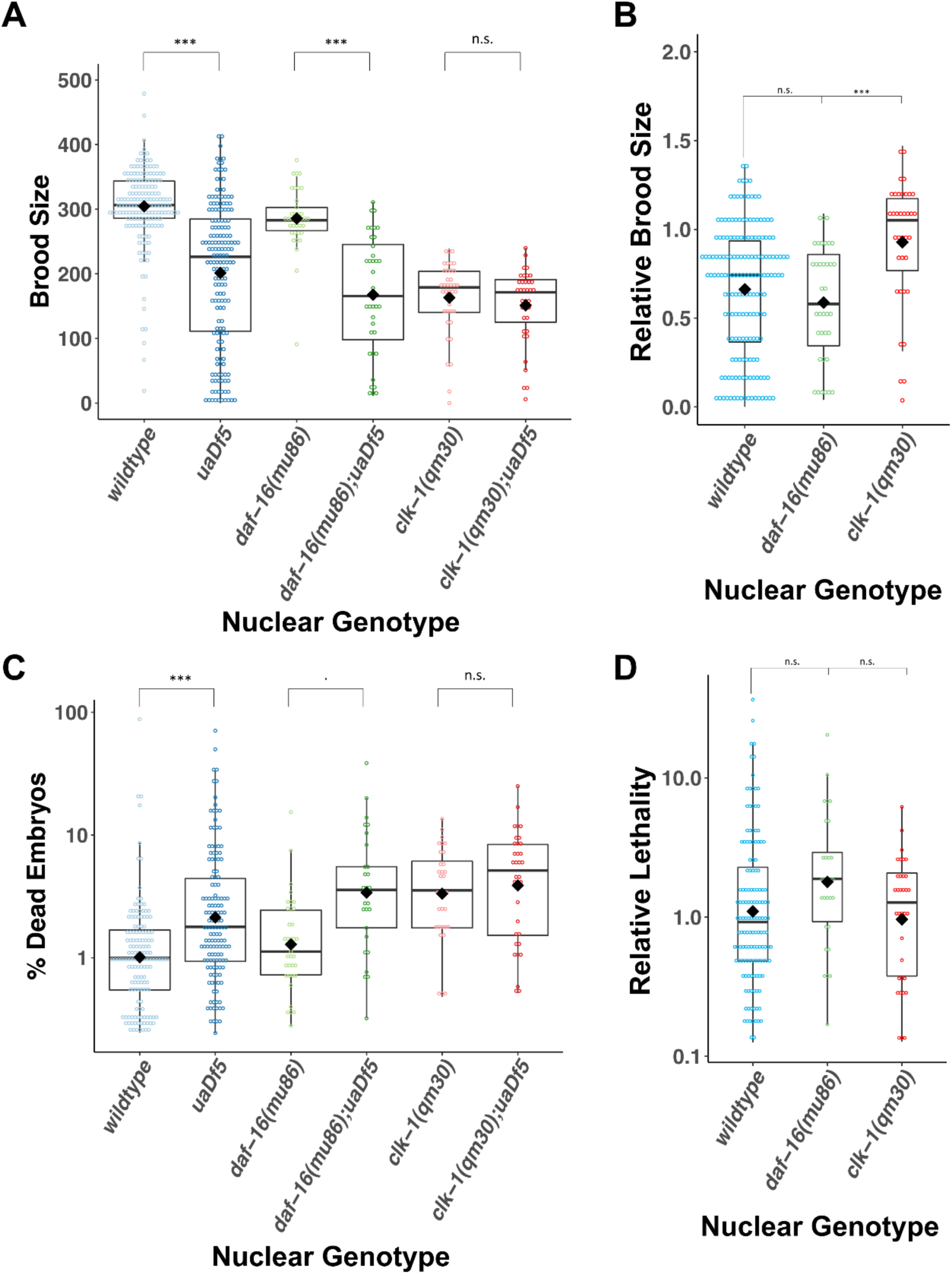
*uaDf5* differentially impacts fitness parameters in lifespan-affecting mutants. **(A-B)** Brood size analysis showing how *uaDf5* differentially impacts brood size in lifespan mutant backgrounds. *uaDf5* has no negative impact on the long-lived mutant *clk-1* but has a modestly larger negative impact on the short-lived mutant *daf-16* than it does in the wildtype background. **(B)** Relative brood size of the animals with and without *uaDf5* in the indicated mutant backgrounds. **(C-D)** Embryonic lethality analysis showing how *uaDf5* differentially impacts embryonic lethality in lifespan mutant backgrounds. *uaDf5* has no negative impact on the long-lived mutant *clk-1* but has a modestly larger negative impact on the short-lived mutant *daf-16* than it does in a wildtype background. **(D)** Relative lethality of the animals with and without *uaDf5* in the indicated mutant backgrounds. Box plots show median and IQR, and the diamond indicates the mean. For **A and C**, statistical analysis was performed using the Mann-Whitney test. For **B and D**, statistical analysis was performed using the Kruskal-Wallis test. (*** p < 0.001, ** p < 0.01, * p < 0.05, . p < 0.1, n.s. not significant).

**Supplementary Table 1:**
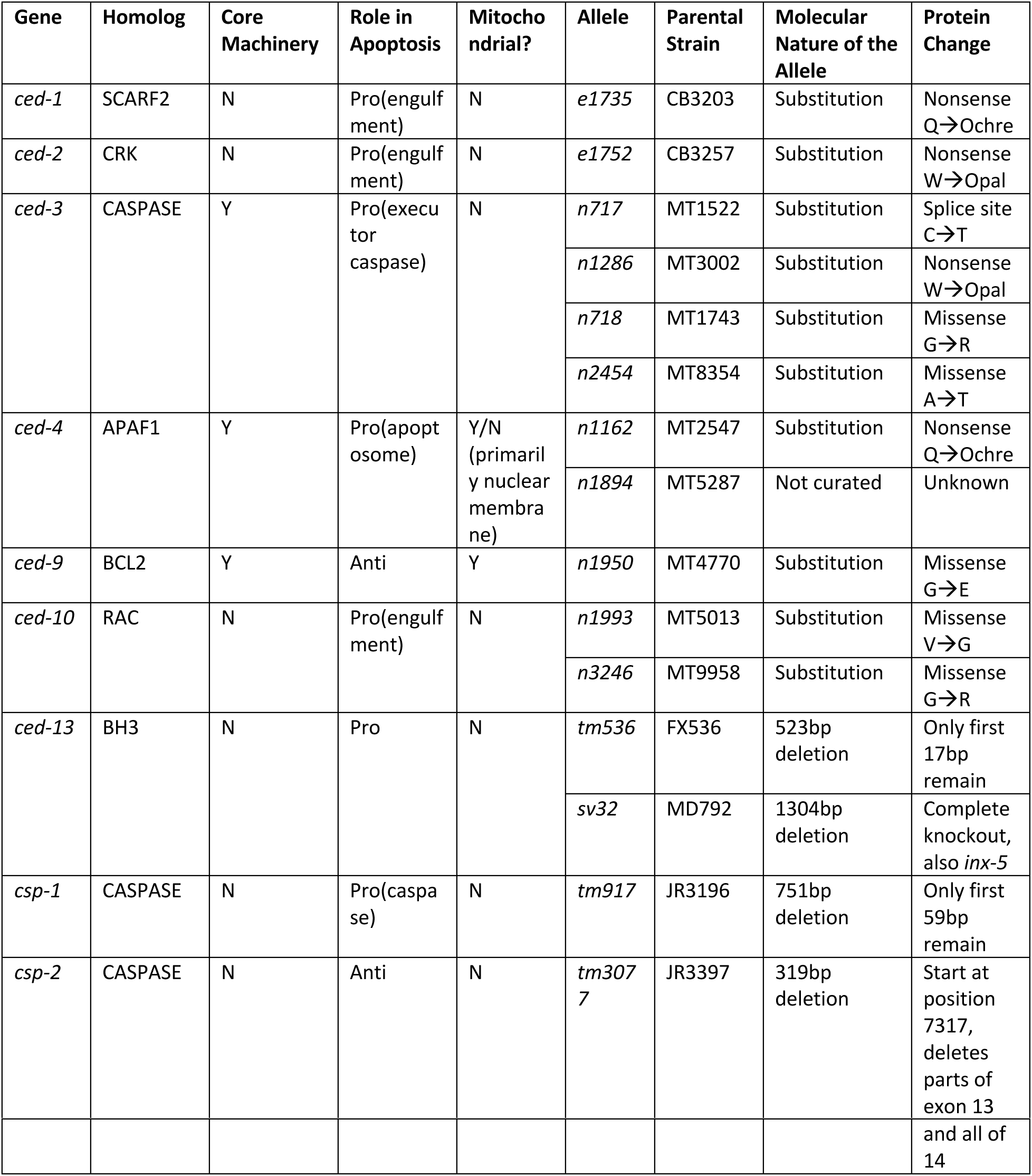
A summary of all mutants analyzed in the PCD pathway, including their known homologs, whether they are part of the core PCD machinery, if they are pro-apoptotic or anti-apoptotic, whether they are mitochondrial proteins, and molecular details of the alleles analyzed.

**Supplementary Table 2:** A summary of all lifespan mutants analyzed, including their known homologs, cellular pathways they are known to act in, whether the mutant extends or reduces lifespan, and molecular details of the alleles analyzed.

| Gene | Homologs | Pathway | Allele | Parental Strain | Molecular Nature of the Allele | Protein Change | Mutant lifespan | Note |
| --- | --- | --- | --- | --- | --- | --- | --- | --- |
| *aak-2* | AMPK | IIS pathway, adult lifespan, peptidyl-serine phosphorylation, regulation of protein localization | *gt33* | TG38 | 606bp deletion | Starts at position 3979, deletes exon 3 | Decrease | - |
|  |  |  | *ok524* | RB754 | 408bp deletion | Starts at position 4136, deletes part of exon 3 | Decrease | - |
| *clk-1* | COQ7 | Adult behavior, regulation of cellular metabolism | *qm30* | MQ130 | 590bp deletion | Starts at position 1044, deletes part of exon 4 and 5 | Increase | - |
| *daf-2* | IGFR | IIS pathway, cellular response to salt, negative regulator of macromolecule metabolic processes, positive regulation of developmental growth | *e1391* | DR1574 | Substitution | Missense P→L | Increase | ts |
|  |  |  | *e1370* | CB1370 | Substitution | Missense P→S | Increase | ts |
|  |  |  | *m41* | DR1564 | Substitution | Missense G→E | Increase | ts |
| *daf-16* | FOXO | IIS pathway, defense response to bacterium, regulation of cellular biosynthetic process, regulation of post- | *mu86* | CF1038 | 1098bp deletion | Deletes 5 exons | Decrease | - |
|  |  |  | *mgDf50* | GR1307 | 20193bp deletion with TCTTCATTTT CAG insertion | Deletes 7 exons | Decrease | - |
|  |  | embryonic development |  |  |  |  |  |  |
| *fem-3* | novel | Male somatic sex determination, masculinization of hermaphrodite germline, positive regulation of macromolecule metabolic process | *q20* | JK816 | Not curated | Unknown | Increase | Gof and ts; female germline development inhibited |
| *glp-4* | VARS | Cell fate specification, embryonic pattern specification, positive regulation of cell proliferation | *bn2* | SS104 | Substitution | Missense G→D | Increase | ts; germline development inhibited |

**Supplementary Table 3:** *C. elegans* strains used in this study.

| Strain | Genotype | Source |
| --- | --- | --- |
| N2 | wildtype | CGC |
| LB138 | *him-8(e1489) IV; uaDf5/+* | CGC |
| JR3630 | *N2; uaDf5/+* (8x backcross version of LB138, *him(e1489)* was outcrossed) | This study |
| JR3688 | *pink-1(tm1779); uaDf5/+* | This study |
| JR3880 | *ced-3(n717) IV; uaDf5/+* | This study |
| JR4330 | *csp-1(tm917) II; uaDf5/+* | This study |
| JR3977 | *ced-3(n1286) IV; uaDf5/+* | This study |
| JR3955 | *ced-13(sv32) X; uaDf5/+* | This study |
| JR3976 | *ced-13(tm536) X; uaDf5/+* | This study |
| JR3926 | *ced-10(n1993) IV; uaDf5/+* | This study |
| JR4001 | *ced-10(n3246) IV; uaDf5/+* | This study |
| JR3972 | *ced-1(e1735) I; uaDf5/+* | This study |
| JR3978 | *ced-2(e1752) IV; uaDf5/+* | This study |
| JR4010 | *ced-1(e1735)I; ced-2(e1752)IV; uaDf5/+* | This study |
| JR3938 | *ced-3(n2454) IV; uaDf5/+* | This study |
| JR3925 | *ced-4(n1162) III; uaDf5/+* | This study |
| JR4017 | *ced-4(n1894) III; uaDf5/+* | This study |
| JR3986 | *csp-2(tm3077) IV; uaDf5/+* | This study |
| JR4027 | *ced-9(n1950) III; uaDf5/+* | This study |
| JR3983 | *ced-3(n718) IV; uaDf5/+* | This study |
| MT1522 | *ced-3(n717) IV* | CGC |
| FX536 | *ced-13(tm536) X* | CGC |
| JR3966 | *glp-4(bn2) I; uaDf5/+* | This study |
| JR3949 | *fem-3(q20) IV; uaDf5/+* | This study |
| JR3937 | *atfs-1(et15) V; uaDf5/+* | This study |
| JR3960 | *daf-2(e1370) III; uaDf5/+* | This study |
| JR3963 | *daf-2(e1391) III; uaDf5/+* | This study |
| JR3993 | *clk-1(qm30) III; uaDf5/+* | This study |
| JR4060 | *daf-2(e1391) clk-1(qm30) III; uaDf5/+* | This study |
| JR3995 | *daf-16(mu86) I; uaDf5/+* | This study |
| JR3997 | *daf-16(mgDf50) I; uaDf5/+* | This study |
| JR4011 | *aak-2(ok524) X; uaDf5/+* | This study |
| JR4012 | *aak-2(gt33) X; uaDf5/+* | This study |
| JR4008 | *daf-16(mu86) I; daf-2(e1391) III; uaDf5/+* | This study |
| JR4005 | *daf-16(mu86) I; daf-2(e1370) III; uaDf5/+* | This study |
| JR3941 | *glp-1(q231) III; uaDf5/+* | This study |
| CF1038 | *daf-16(mu86) I* | CGC |
| MQ130 | *clk-1(qm30) III* | CGC |
| JR3981 | *csp-1(tm917) II; ced-3(n717) IV; uaDf5/+* | This study |

## Acknowledgments

We would like to thank the members of the Rothman lab for their support. We thank Jen Smith and the BNL at UCSB for providing excellent facilities which were necessary for this work. We thank Kyle Ploense for his help in statistical analysis. Worm strains used in this work were provided by the Mitani lab, as well as the Caenorhabditis Genetics Center (CGC), which is funded by NIH Office of Research Infrastructure Programs Grant P40 OD010440.

## Author contributions

Sagen E. Flowers - conceptualization, investigation, funding acquisition, writing – original draft preparation, writing – review and editing

Rushali Kothari – investigation, writing – review and editing

Yamila N. Torres Cleuren - investigation, writing – review and editing

Melissa R. Alcorn – statistics, writing – review and editing

Chee Kiang Ewe - writing – review and editing

Geneva Alok - writing – review and editing

Pradeep M. Joshi – conceptualization, investigation, writing – review and editing

Joel H. Rothman - conceptualization, supervision, funding acquisition, writing – review and editing

## Data Availability

All data generated or analyzed during this study are included in this article and accompanied supplementary materials. The raw reads of the sequenced N2, LB138, and JR3688 genomes have been deposited at the NCBI SRA under the study accession number PRJNA836592.

